# Neural Correlates of High-Level Visual Saliency Models

**DOI:** 10.1101/2023.07.29.551075

**Authors:** Alexander Kroner, Mario Senden, Rainer Goebel

## Abstract

Visual saliency highlights regions in a scene that are most relevant to an observer. The process by which a saliency map is formed has been a crucial subject of investigation in both machine vision and neuroscience. Deep learning-based approaches incorporate high-level information and have achieved accurate predictions of eye movement patterns, the overt behavioral analogue of a saliency map. As such, they may constitute a suitable surrogate of cortical saliency computations. In this study, we leveraged recent advances in computational saliency modeling and the Natural Scenes Dataset (NSD) to examine the relationship between model-based representations and the brain. Our aim was to uncover the neural correlates of high-level saliency and compare them with low-level saliency as well as emergent features from neural networks trained on different tasks. The results identified hV4 as a key region for saliency computations, informed by semantic processing in ventral visual areas. During natural scene viewing, hV4 appears to serve a transformative role linking low- and high-level features to attentional selection. Moreover, we observed spatial biases in ventral and parietal areas for saliency-based receptive fields, shedding light on the interplay between attention and oculomotor behavior.

## 1. Introduction

The study of saliency as an underlying representation that guides eye movement behavior in humans has garnered interest from both the machine vision and neuroscience communities. In neuroscience, it has been formalized as a pre-attentive map that captures the relative conspicuity of spatial locations in the visual field and guides shifts of attention [1]. Computational models have attempted to provide a mechanistic account of feature extraction and integration to mimic the neural processes that give rise to a saliency map. One influential example by Itti et al. [2] derives saliency maps on the basis of orientation, intensity, and color contrasts. This early model and similar low-level approaches were capable of explaining fixation patterns better than chance. Their popularity was, moreover, grounded in the concordance between model mechanisms and our knowledge of early visual processing. However, low-level models were later deemed insufficient to accurately capture fixation behavior in response to natural scenes [3, 4, 5]. A shift in understanding towards the dominant role of object information for attentional selection [6, 7, 8], as already suggested in the seminal eye movement experiments by Yarbus [9], has then paved the way for the successful application of deep neural networks. These have since proven to be highly accurate for saliency map predictions [10], thanks to their ability of detecting high-level features in complex images. It must be noted that deep saliency models also leverage low-level and, to a lesser degree, mid-level features [11]. As a consequence, the dichotomy between low- and high-level approaches can be considered simplified but nevertheless allows us to investigate which feature basis is more suitable for explaining neural activity after careful disentanglement of their respective contributions.

While computational work has made substantial strides towards building accurate models of saliency, the question remains whether a topographical representation of saliency even exists in the brain and if so, where it is located [12]. Previous studies have suggested neural correlates of saliency in areas such as the superior colliculus (SC) [13], pulvinar [14], visual cortices V1 [15] and V4 [16], the temporal [17] and parietal lobe [18], as well as the frontal eye fields (FEF) [19]. Many of these observations resulted from experiments that either elicited a response of saliency through simplistic stimuli, such as an array of targets and distractors distinguished by color or form, or defined saliency maps in terms of low-level visual features. Although the intraparietal sulcus (IPS) [20], FEF, and SC have repeatedly been found to partake in oculomotor planning, it is unlikely that they alone give rise to naturalistic viewing behavior [16], which is influenced by semantic information such as face and text detection [21]. Their close ties to the oculomotor system rather support fast orienting responses that are necessary for adapting to sudden changes in the environment [22]. Instead, ventral visual areas may contribute more semantic aspects to the allocation of attention through coordinated neural interactions with frontoparietal structures. Mazer and Gallant [16], for instance, consider V4 a central hub for computing the locations of salient features and the formation of a stimulus-driven saliency map that is forwarded to down-stream areas. Anatomical projections to the frontoparietal lobe [23] could thereby mediate the influence of high-level scene analyses on saccadic control for complex visual inputs.

To systematically address the question of whether saliency maps are indeed represented in the brain, several criteria must be fulfilled [12]. First, the activity levels of neurons that encode saliency reflect the degree to which an image region is salient regardless of the specific features that render it salient. Second, a cortical saliency map is required to be topographically organized, such that a selection mechanism in the oculomotor system can determine the next fixation target and produce saccadic eye movements. Third, the locus in question must hence be functionally connected to the oculomotor system. Lastly, the saliency map captures high-level spatial features to serve as an underlying representation of eye movements for naturalistic stimuli. These characteristics do not rule out a distributed notion of attentional processing, where aspects other than feature-driven responses, such as the integration with top-down signals towards the formation of a priority map [25], occur elsewhere.

Paralleling the shift in computational modeling towards more naturalistic stimuli, we wanted to examine the neural underpinnings of high-level saliency using models that have demonstrated compelling predictive performance with respect to fixation patterns. We hypothesize that ventral visual processing is necessarily involved through its spatial encoding of objects in vivo, akin to deep classification networks in silico [26]. To test this assumption, a link between algorithmic representations and neural responses needs to be established. Encoding models can create this link by making predictions about the neural response within a specific region at either the level of individual measurement channels or their multivariate patterns of activity, based on the stimulus descriptions of a candidate model. Applying this method to functional magnetic resonance imaging (fMRI) data has become an important tool for evaluating model predictions in terms of their biological plausibility. Likewise, it allows to query whether certain information is present in brain activity [27], especially when measured at higher field strengths [28]. Here, we assume that if saliency maps are indeed the underlying representation for the selection of fixation targets, then we can detect their neural signature even in the absence of eye movements. This assumption is corroborated by experiments that have successfully decoded image saliency from fMRI activity recorded during central fixation [29, 30].

In our study, the aim was to shed light on the relation between model-based saliency, as a proxy for the spatial distribution of real eye movements, and the brain. We hence formulated the following questions: (1) Are low- and high-level saliency maps differentially represented in the brain? (2) How do neural predictions from high-level saliency models compare to deep neural networks trained on different tasks? (3) Are saliency maps topographically organized after disentangling their contribution from general visual features? To address these questions, we applied several methods that link model representations to the brain and contrasted the predictions from different model outputs or layer activations (see Figure 1). This approach is aimed at localizing the cortical regions involved in saliency computations and characterize their functional role subserving naturalistic scene processing. Our results were remarkably consistent across methodologies and suggest that high-level saliency models encapsulate information that is highly relevant with respect to the cortical visual system. Parietal and especially ventral visual areas were found to be critically involved in saliency computations, with the midventral *region of interest* (ROI), which corresponds to hV4, playing a central and integrative role. As a result of our primary analysis, we also noted a spatial bias of receptive fields related to visuospatial attention towards the upper and left visual field with possible effects on oculomotor behavior. This constitutes the first instance of linking model-based semantic saliency maps to a large-scale fMRI dataset of complex scenes to the best of our knowledge. Our findings can thus contribute to a deeper understanding of the neural and computational factors that underlie naturalistic free-viewing behavior.

**Figure 1:**
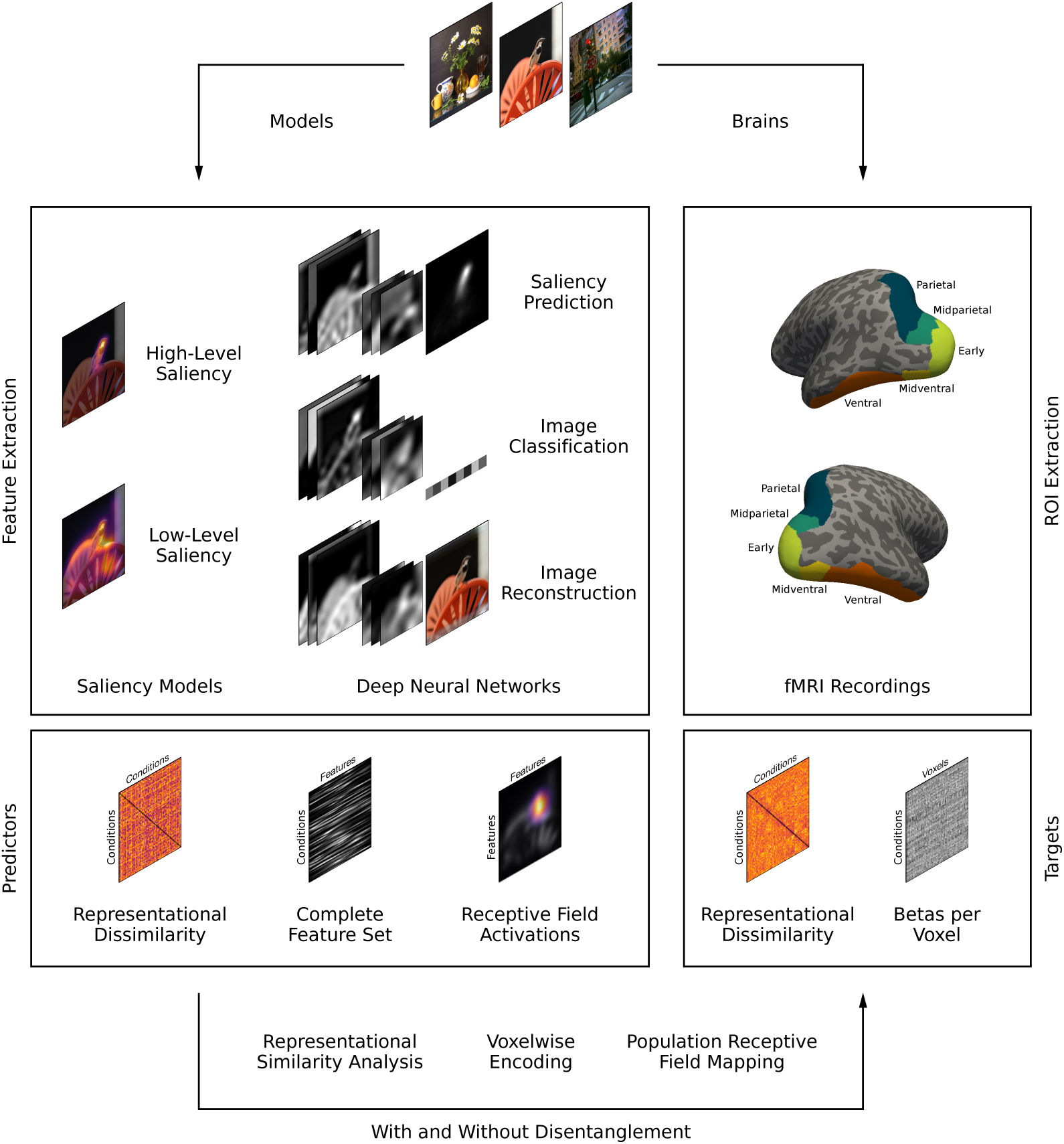
A summary of our approach towards investigating the neural correlates of high-level saliency. Complex images were presented to participants in the *Natural Scenes Dataset* (NSD) [24] and formed the input to low- and high-level saliency models as well as deep neural networks trained on different tasks. To establish a direct link between models and neuroimaging data, multiple techniques were applied that each require a specific representation format of both the predictors and targets. Feature spaces consisting of saliency maps or layer activations were either disentangled or treated separately to quantify and contrast the predictive accuracy of high-level saliency.

## 2. Materials and Methods

### 2.1. Natural Scenes Dataset

In this study, we used the *Natural Scenes Dataset* (NSD) [24]. It comprises large-scale 7 T fMRI data acquired from eight participants with a whole-brain coverage at 1.8 mm resolution. Each participant viewed 9,000–10,000 unique images of natural scenes from the Microsoft *Common Objects in Context* (COCO) database [31]. Images were presented up to three times with a size of 8.4° × 8.4°, resulting in 22,000–30,000 trials distributed over 30–40 scan sessions within one year. Throughout the experiment, participants were instructed to fixate centrally and perform a long-term recognition task in which they had to indicate whether the current image had previously been presented in any of the sessions. Measurements of fMRI responses were made available in both volume- and surface-based versions. Further details about NSD can be found in the publication by Allen et al. [24].

### 2.2. Dataset Preparation

We utilized the 1.8 mm volume preparation of the NSD in which the *hemodynamic response function* (HRF) was estimated per voxel, an adaptation of *GLMdenoise* [32] was applied for denoising, and ridge regression was fitted to better estimate single-trial betas. The resulting betas were normalized across trials within each scan session via z-scoring to remove unwanted variability over time. All trials were concatenated per participant, excluding the three scan sessions that were held out by the NSD authors for benchmarking purposes, and averaged over repeated presentations (1–3 times) of the same image. We extracted visually evoked responses within five ROIs defined by the provided *streams* delineation that encompasses neural activations along the dorsal and ventral visual pathways: *early*, *midventral*, *ventral*, *midparietal*, and *parietal* (see Table A.1 for details). Unless otherwise stated, analyses were performed on a set of 5,000 unique conditions per subject.

### 2.3. Surface Visualizations

For surface-based visualizations, we transformed our analysis results to the native space of individual subjects. The volume-based data was first mapped to three cortical depth surfaces at 25%, 50%, and 75% of the distance between the pial and white matter surface using linear interpolation, and averaged across the depth projections. Results were then spatially smoothed using a geodesic Gaussian kernel (FWHM = 3 mm) and normalization procedure based on the vertex area. Finally, they were mapped to the *fsaverage* brain space via nearest-neighbor interpolation and averaged across subjects for each vertex if data from at least five out of eight subjects was available. All quantities intended for surface visualizations were computed from the results only after these transformations to avoid incorrect averaging or smoothing of data formats, such as circular values. Group-average data was visualized with respect to the streams ROIs, where all analysis steps were performed, but also a more detailed view of retinotopically defined areas V1, V2, V3, and hV4. For this purpose, the fsaverage surface was projected to a flatmap in the former case and a sphere in the latter case. Areas V1, V2, and V3 hereby corresponded to the early ROI and hV4 to the midventral ROI.

### 2.4. Saliency Models

To generate continuous saliency maps from images of natural scenes and contrast the differential contribution of feature complexity, we selected four low-level models (AIM [33], GBVS [34], ICF [35], IKN [2]) and four high-level models (DVAP [36], MSI [37], SalGAN [38], SAM [39]) for comparison. A summary of these models’ results on the MIT/Tübingen saliency benchmark [40] can be seen in Table B.2 and their predictions on a set of example images in Figure C.11. All models, except for our own network (MSI), were available via the *SMILER* [41] software package and tested with their default settings. Unique images for each participant were then provided as input to the models at a downsampled resolution of 320 × 320 px using the area interpolation method. Instances from the COCO database that were either part of the training or validation set used to tune the high-level saliency models were excluded from further analysis to avoid potential data leakage. This amounts to approximately 15% fewer images per participant.

#### Low-Level Models

Although unified by the common task of saliency prediction, low-level models employed a diverse range of methodologies. They generally took inspiration from biological mechanisms and computed early visual features akin to neural processing of local features and center-surround differences. However, the design of such model architectures was not only driven by theory but also the computational analysis of image data. **AIM** [33] determines saliency based on the measure of self-information across image regions. To do so, a set of local and independent features is first extracted from random image patches of natural scenes and forms the basis for a statistical distribution. The saliency within each region is then inversely related to this likelihood. **IKN** [2] constructs a set of filters for colors, intensities, and orientations at multiple spatial scales and calculates center-surround differences within each channel. Therefore, the algorithm detects feature contrast rather than feature strength in images. The resulting maps are normalized and combined into a common representation of saliency. **GBVS** [34] utilizes the same underlying feature set but instead interprets the positions in all feature maps as the nodes of a fully-connected graph. The weight of edges is proportional to the proximity between nodes and the dissimilarity of their values. This culminates in a definition of saliency, where differences between surrounding nodes result in higher scores. **ICF** [35] learns a point-wise nonlinear combination of features derived from local intensity and contrast in natural images. Once again, the feature space is spanned by multiple spatial scales using a Gaussian pyramid. The combined output is then blurred and a center bias added to the prediction of saliency maps.

#### High-Level Models

To ensure comparability between different deep learning methods, only models with a VGG16 backbone [42] pre-trained on the ILSVRC dataset [43] for object classification were included in this study. We used the version of each model where weights were tuned towards saliency prediction on the SALICON dataset [44]. Therefore, any observed differences between the models are primarily the result of architectural decisions and optimization specifics. Here we summarize the former. **DVAP** [36] extracts activations from three mid- to high-level stages of the VGG16 encoder to incorporate features with different complexity and receptive field sizes. Each of these outputs is trained to individually predict saliency maps using deconvolutional layers, as well as through a combined multi-scale representation. **MSI** [37] augments object-based representations with multi-scale information and a map of global image properties towards contextual processing of spatial features. This approach emphasizes the importance of treating objects as embedded in a natural scene rather than in isolation. To that end, mid- to high-level activations are sampled at multiple receptive field sizes using an *Atrous Spatial Pyramid Pooling* (ASPP) module [45]. **SalGAN** [38] consists of a generator and a discriminator that together form a *generative adversarial network* (GAN). The former produces saliency maps given a natural image whereas the latter learns to distinguish between generated and ground truth saliency maps. **SAM** [39] leverages a convolutional version of *long short-term memory* (LSTM) units that operate along the spatial dimensions of a high-level feature tensor. Additionally, the model incorporates a spatial attention mechanism to selectively focus on the regions of this tensor that are most relevant for saliency prediction.

### 2.5. Feature Extraction

All saliency maps generated by the aforementioned list of models were saved at a resolution of 128 × 128 px for further analysis. Moreover, we extracted activation maps from several convolutional layers of the MSI network (see Figure 2). A total of 17 trained layers was selected, of which the first 13 were initialized from pre-trained VGG16 weights and the remaining ones according to the *Glorot Uniform* method [46]. We can therefore compare how fine-tuning the network towards saliency prediction changes the representations of its layers. Consequently, we also extracted features from the VGG16 network trained on ILSVRC for object classification. All activation maps were spatially resized via the area interpolation method while keeping the number of channels fixed, such that the resulting tensor (*height* × *width* × *channels*) consists of approximately 16,000 elements. The input sizes to MSI and VGG16 were 320 × 320 px and 224 × 224 px respectively.

**Figure 2:**
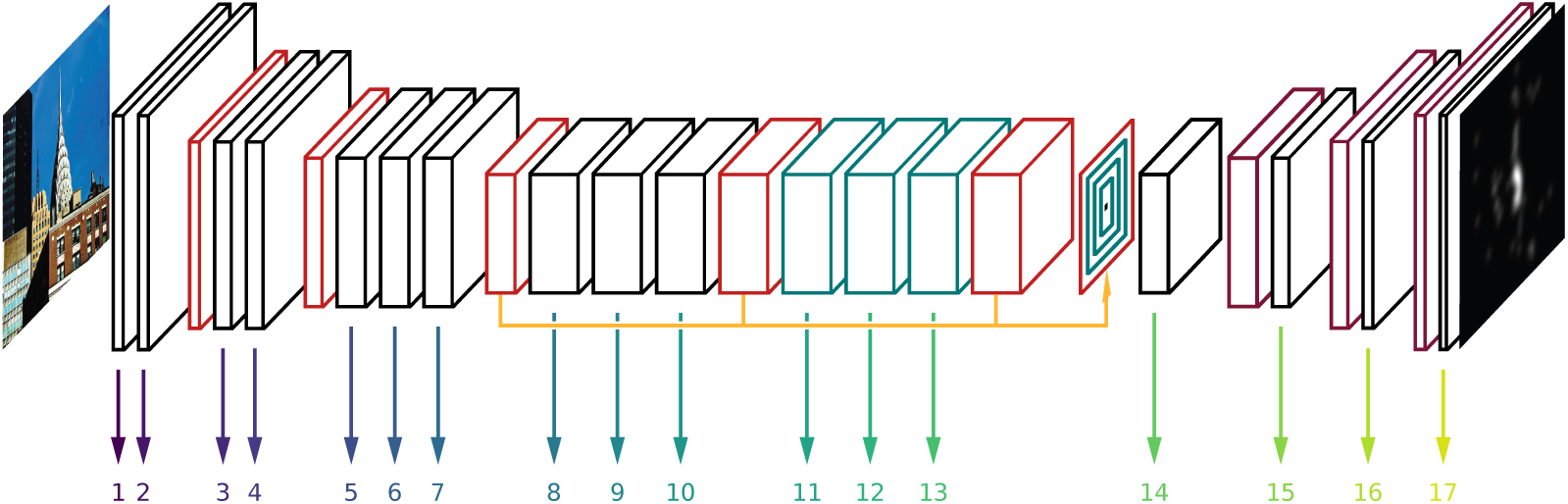
The encoder-decoder architecture of the MSI network with arrows indicating extracted layer outputs. **Black** boxes represent convolutional layers and **cyan** boxes convolutional operations with a dilation factor larger than 1. **Red** and **purple** boxes denote down- and up-sampling layers respectively. The **yellow** arrow visualizes skip connections that feed into a multi-scale ASPP module. Note that numbers reflect the *n*th feature space extracted for subsequent analyses and not the *n*th processing layer of the network. Resizing layers and intermediate outputs from the ASPP module were not included here. This figure is adapted from Kroner et al. [37].

Notably, the MSI and VGG16 networks not only differ by training task but also their architecture beyond layer 13. The former implements a fully convolutional decoder that preserves and transforms the spatial information of feature maps, whereas the latter combines all information through location-independent dense layers towards a class label. To infer whether any observed differences are due to changes in either the task or architecture, we trained an autoencoder with the identical architecture as MSI on stimulus reconstruction rather than saliency prediction. This alternative model is trained for 100 epochs on the same set of stimuli as MSI and minimizes the mean squared error between reconstructed and original RGB images. Finally, its layer activations were extracted from a model snapshot with the best performance on a held-out validation set.

### 2.6. Center Bias Removal

Fixation patterns under experimental settings are subject to a spatial bias, where regions close to the image center are preferentially targeted independent from the distribution of features [47]. Such a prior has since been captured explicitly (ICF, SAM) or occurred implicitly (AIM, GBVS, IKN) [48] in saliency maps generated by computational models. To distinguish the contribution of feature information from this prior, we removed the center bias in two stages. First, all values of either saliency maps or layer activations were z-scored for each image independently. Second, we applied the QuantileTransformer function of *scikit-learn* [49] to transform pixel values based on a rank order procedure, similar to the approach put forth by Rahman and Bruce [50]. To that end, 25 evenly spaced quantiles between 0 and 1 were computed for a given pixel’s values across all conditions. An estimated cumulative distribution function then served to map original values to their rank, producing a uniform output distribution between 0 and 1. This was repeated for all pixel locations independently. The effect of this two-step procedure on the center bias is illustrated in Figure D.12.

### 2.7. Representational Similarity Analysis

To link the activation patterns of saliency maps and layer outputs to the brain, we performed *representational similarity analysis* (RSA) [51]. This method estimates the representational geometry of a given system by comparing its responses for each pair of conditions [52]. For that purpose, we derived *representational dissimilarity matrices* (RDMs) from model and brain activations respectively across the stimuli. Pairwise distances *d* between the multivariate response patterns were computed according to the Pearson correlation coefficient ρ via *d* = 1 − ρ. To infer which model best accounts for the neural data, we related the RDMs by means of a second-level similarity analysis. Here, we chose Pearson correlation as the summary statistic, assuming that the ratio scale of dissimilarities in the model RDMs, and not only the rank order, corresponds to neural RDMs [53]. These calculations were limited to the upper triangle of the symmetric RDMs.

#### Feature Space Grouping

We further used RSA to investigate the differential contribution of low- and high-level models by grouping all saliency map RDMs into the respective low- or high-level category. In practice, we averaged the representations within each class [54] and thus obtained two sets that represent a dichotomy of methods towards saliency prediction. To decorrelate the approaches and disentangle their similarity with neural responses, we performed a *zero component analysis* (ZCA) whitening procedure, similar to Greene and Hansen [55]. This technique defines a symmetric whitening matrix as *W* = Σ^−1/2^, where Σ denotes the covariance matrix of the inputs. After applying the whitening matrix to the RDMs, the covariance becomes equal to the identity matrix. This effectively decorrelates the two spaces but maintains a high correlation between original and whitened features, which largely preserves their initial meaning [56]. The grouped results for low- and high-level saliency models then served as the basis for a comparative analysis.

#### Univariate Noise Normalization

It must be noted that the reliability of neural RDMs is reduced by the measurement noise that affects each voxel [57]. Although a multivariate noise normalization procedure was proposed to counteract this issue [57], its benefits across different fMRI datasets remain inconclusive [58]. The GLMdenoise procedure, which was used to reliably estimate beta values of NSD, has however demonstrated robust improvements regarding the consistency of RSA results [58, 59]. We thus only combined it with a simpler univariate noise normalization approach and divided each voxel value by its variance. The variance for each voxel and subject was calculated as the mean of variances across conditions where all three repetitions were available. This effectively reduces the contribution of noisy voxels and converts betas to t-values.

#### Noise Ceiling Computation

The noise ceiling of correlation results between model RDMs and neural RDMs was defined based on a lower and upper bound. They were estimated by the Pearson correlation between each single-subject RDM with the group-average RDM that either excludes (lower bound) or includes (upper bound) the subject in question [60]. The upper bound hence overestimates the achievable model performance given the measurement noise and variability across subjects. Before correlation values were averaged across subjects, we converted them using the Fisher z-transformation. The final noise ceiling results were then reverted to Pearson correlation coefficients.

### 2.8. Voxelwise Encoding Models

Voxelwise encoding is another popular approach to test hypotheses generated by computational models regarding the neural responses for a set of stimuli. It allows the study of how well the representations in a candidate model are able to explain representations in the brain. In practice, encoding models map features to individual voxels by fitting the weights of a linear regression and thus learn to predict the neural activity for novel stimuli. Unlike RSA, this method characterizes single voxel responses rather than populations thereof and is suitable for examining the spatial distribution of features on the cortical surface [61]. Here, we applied two techniques: regularized linear regression and *population receptive field* (pRF) mapping. The performance of a candidate model was quantified via the median Pearson correlation between predicted and true beta values across all voxels [62].

#### Ridge Regression Pipeline

Ridge regression is a common choice for establishing a direct connection between candidate models and neural responses due to its simplicity [63, 64]. In this study, we also make use of this technique to test whether certain features are explicitly represented in the brain and to perform a model comparison [65]. The feature space consists of saliency maps or layer activations and the targets are all voxels within an ROI for one subject at a time. Before fitting the weights of an encoding model, we standardized the input features across conditions. Linear regression with L2-regularization was selected to reduce the risk of overfitting and improve its generalization performance. Its regularization strength was optimized by searching over eight log-spaced shrinkage values between 10^1^ and 10^8^. One optimal value was found per voxel by minimizing the least-square error on a held-out validation set comprising 20% of all training data. Taken together, this approach combines feature responses independent of their spatial position to predict neural responses for each voxel. Hereafter, we refer to this technique as ENC.

#### Topographical Analysis

An alternative encoding technique is pRF mapping, which retains a correspondence between the spatial arrangement of information in the visual space and voxel activity in the brain space. In practice, the location and size of receptive fields, assuming an isotropic Gaussian shape, are optimized with respect to the neural betas based on a grid-search procedure [66]. This can give rise to a systematic investigation of eccentricity and polar angle preferences on cortical surfaces. While this mapping is commonly performed for simple stimulus descriptions, such as a binary map of pixels that either lie inside or outside the stimulus shape, it can also be based on feature maps from convolutional neural network layers [67]. This instead tests whether complex feature representations are topographically organized in the visual system.

Here, we adopted the *feature-weighted receptive field* model by St-Yves and Naselaris [67] to fit saliency maps or layers to individual voxels. The search space for optimal pRF parameters is spanned by 10 log-spaced receptive field sizes σ between 0.3 and 4.0, where locations are linearly sampled with a distance of 1 σ from each other. We also included locations outside the field of view (up to a factor of 1.1), since such receptive fields can still be reliably mapped when using a grid-search procedure [68]. This yields a total search space of 2,280 pRF parameter combinations. After filtering the feature maps with the candidate receptive fields, we perform ridge regression for each voxel within a region of interest as described before. The combination of receptive field size and location that best describes a voxel’s response across conditions, according to a held-out validation set (20% of all training data), was then retained for further analysis. Final predictions were generated based on coefficients that were re-fitted on the entire set of training and validation data.

#### Feature Space Grouping

To accurately predict neural activity in a region of interest, it has been shown that combining features from different stages of the processing hierarchy in deep neural networks can outperform single layers [67, 69]. We thus fitted whole networks onto neural betas by using the aforementioned ridge regression pipeline. To that end, the chosen set of layer activations was concatenated and served as the joint predictors of an encoding model. This can generate insights as to how closely the distributed representations learned throughout an entire model resemble cortical information. By contrasting the predictive performance of models with the same architecture, we can further evaluate the impact of the training task on this measure. The grouping of saliency maps, generated by either low- or high-level models, followed the same procedure. In contrast to RSA, we thus stacked rather than averaged the representations within each class.

Combining multiple feature spaces into a joint model allows us to capture complementary information towards the prediction of neural responses. However, standard ridge regression only optimizes a single regularization parameter for all spaces, which ignores their specific characteristics. To address this problem, *banded ridge regression* [70] is an extension that fits a separate parameter for each space. This can improve the prediction accuracy of a joint model but also disentangle the contribution of individual feature spaces by reducing spurious correlations between them [70, 71]. The performance of a feature space is then the result of splitting the joint model’s correlation score between predicted and true neural activity into a sum of individual components. Hence, this approach facilitates a meaningful comparison of models or features and their interpretation. We performed all computations of banded ridge regression and scoring methods via the *himalaya* package in Python [71].

In the case of pRF mapping, we applied the same technique to examine whether the unique contribution of a specific feature, such as saliency maps, gives rise to a topographic organization in the brain. To that end, all feature spaces were first filtered by the same candidate receptive field and then mapped to beta values via banded ridge regression. This yielded a predictive score per feature space and was repeated for all pRF parameter combinations to find the best fit per voxel. One assumption of this procedure is that spurious correlations between feature spaces primarily exist for the same receptive field. If, however, the topographic organization of feature spaces is considerably different, then the proposed method may not sufficiently disentangle their separate contribution to the joint encoding model. This would instead require the application of banded ridge regression to feature spaces with individual pRF parameters, which would render the search space computationally intractable.

#### Noise Ceiling Computation

Although we are mostly interested in the relative ranking between different features or models, the absolute performance of an encoding method is informative of its ability to capture representations in the brain. Predictive accuracy is, however, limited by measurement noise, which can be quantified via the noise ceiling for each voxel. NSD provides the *noise ceiling signalto-noise ratio* (ncsnr) based on a closed-form procedure by Lage-Castellanos et al. [72]. We recomputed this quantity on all data available to the public, i.e. excluding the three held-out scan sessions for each subject. By taking into account the different number of trials used for averaging betas per condition, it can then be transformed to a noise ceiling as described in the NSD manual:

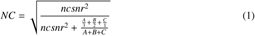

Here, *A*, *B*, and *C* represent the number of conditions based on 3, 2, and 1 trial(s) respectively. We thus obtained an upper bound for the correlation between predicted and true betas per voxel.

### 2.9. Statistical Testing

All computations using the aforementioned methods were carried out with respect to a fivefold cross-validation procedure. This includes the pre-processing of features, fitting of regression weights where applicable, and evaluation of scores and parameters. To obtain robust estimates of model performance and parameters, test set results were averaged across folds. These quantities were then used for statistical testing. Correlation coefficients were transformed using the Fisher z-transformation prior to such tests.

The significance of model results was assessed by calculating the non-parametric Wilcoxon signed-rank test. We performed a one-sided evaluation of whether population scores across the eight subjects are significantly greater than zero. As a basis for this statistical analysis, we quantified the correlation between model and neural RDMs (RSA) or the median correlation between predicted and true voxel activations (ENC, PRF) for each subject and ROI. These quantities were then converted using the Fisher z-transformation. The Wilcoxon test returned p-values that were adjusted using the Benjamini-Hochberg procedure [73] to control the *false discovery rate* (FDR) at a level of 0.05 across models and ROIs. Corrected p-values smaller than 0.05 were considered significant. To evaluate the significance of pairwise model differences, we instead performed a two-sided Wilcoxon test with two paired samples but otherwise followed the same steps.

This statistical testing procedure was also carried out to infer whether the performance of a model category (i.e. low-level or high-level) was significantly greater in one compared to another ROI. The results for each subject and ROI were averaged across the three methods and a onesided Wilcoxon test was performed as before. To assess significance of equivalence between two model categories within one ROI, we applied the *two one-sided tests* (TOST) procedure [74]. It evaluates whether the difference in performance falls into an interval, determined by a lower bound Δ*_L_* and upper bound Δ*_U_*, that is sufficiently small to be considered negligible and thus absent of an effect. Here, the bounds were estimated based on a bootstrapping procedure, where two samples of eight subjects (with replacement) were created for a given model class. This was repeated 1,000 times to obtain a distribution of mean differences between each of the two sets. We then defined both the lower and upper bound as the 99th percentile of this distribution and performed two null hypothesis tests: *H*0_1_ : Δ < −Δ*_L_* and *H*0_2_ : Δ > Δ*_U_* . If both hypotheses are rejected, then we conclude that no significant performance difference between two model classes was found within a given ROI.

Significantly predicted voxels were determined with respect to a non-parametric permutation test for each model, subject, and ROI. Here, we randomly shuffled the conditions of the true beta values, calculated the correlation coefficients with predicted beta values, and repeated this procedure 1,000 times to obtain a null distribution of scores per voxel. P-values were then computed based on (*n* + 1)/(*m* + 1), where *n* is the number of values from the null distribution larger than the observed similarity and *m* the number of permutations [75]. These results were corrected for multiple comparisons across all models, subjects, and ROIs with an FDR level of 0.05. Voxels with a corrected p-value smaller than 0.05 were considered significantly predicted by the model.

We also compared two models regarding the percentage of voxels that are better predicted by one compared to the other. This quantity is defined as the model advantage with values between 0 and 1. To that end, we first calculated the advantage of a given model based on correlation scores for voxels that are significantly predicted by at least one of the two models. A null distribution was then formed after randomly permuting the correlation scores from the two models for each voxel individually and recomputing the advantage for the same model 1,000 times per ROI. If the percentage of voxels that are better predicted by this model is larger than 0.5, then *n* is the number of elements from the null distribution larger than the observed value. Likewise, if the true model advantage is smaller than 0.5, then *n* is the number of elements from the null distribution smaller than the observed value. This was aimed at testing the deviation from parity between the two models. Finally, the same steps for p-value computations, FDR correction, and significance thresholding were performed as before to statistically assess the model differences [67].

## 3. Results

### 3.1. Inter- and Intra-Model Similarities

A subset of 10,000 images was randomly selected from all stimuli presented to the eight participants. They were forwarded to the aforementioned saliency, classification, and autoencoder models to evaluate the similarity across (a) saliency maps of different models and (b) layer activations within a given model. Figures 3A-D visualize the correlation results for eight saliency models and the 18 layers of MSI, whose last layer is equivalent to the predicted saliency map. The two methods for assessing pairwise similarity yielded results with a strong qualitative agreement but correlation values for the elementwise encoding model were generally higher due to the fitting of regression weights. In the case of saliency models (see Figures 3A and 3B), high-level approaches formed a tight cluster whereas low-level models were more loosely related. Both categories demonstrated higher correlation values within than across their respective clusters. This finding was somewhat unsurprising given that all high-level models are deep neural networks that were trained on the same dataset. Nevertheless, these results reinforce the suggested dichotomy between low- and high-level approaches.

**Figure 3:**
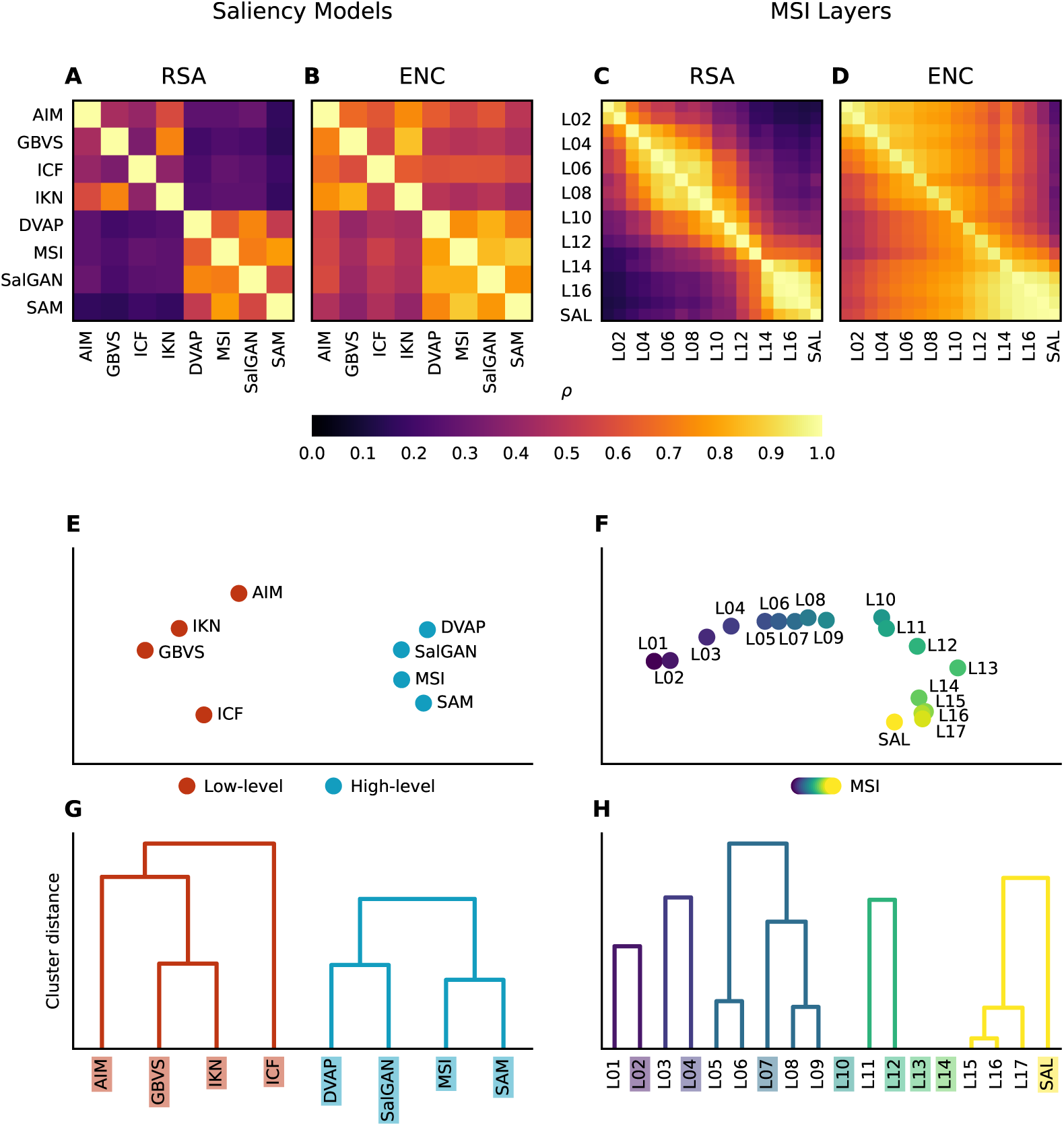
Pairwise comparisons to determine the similarities between saliency maps from eight models and layer activations from MSI specifically. Both the inputs and desired outputs of the analyses were based on features alone and thus unrelated to neural data. (**A** - **D**) Pearson correlation scores ρ were obtained from two methods: RSA (**A**, **B**) and an encoding model (**C**, **D**). Note that the former yielded symmetrical results along the diagonal whereas the latter introduced a directionality from predictors (x-axis) to targets (y-axis). For ridge regression, elementwise correlation results were averaged across saliency maps or layer activations. (**E** - **H**) An analysis of the previous similarity results obtained from the RSA method. Two techniques were employed to visualize and cluster the pairwise correlation scores: MDS (**E**, **F**) and Agglomerative Clustering (**G**, **H**). The former embeds the similarities between models or layers in a two-dimensional space whereas the latter forms a dendrogram that represents a hierarchy of clusters. The height at which two clusters merged in the dendrogram indicated the maximum distance between all contained elements. Note that leaf elements without an associated dendrogram formed their own cluster. Colored labels (**G**, **H**) denote which models or layers were kept for further analyses.

Regarding the similarity between layers within the MSI network (see Figures 3C and 3D), one should keep in mind that the first 13 layers can be viewed as the encoder, with feature weights initialized from VGG16 before fine-tuning, while the remaining layers form the decoder. This division was also reflected by the pairwise correlation results. Decoder representations were highly related to each other but distinct from the encoder layers. Furthermore, the transition layer (L13) between the encoder and decoder cannot easily be assigned to either of the two clusters. A notable asymmetry regarding the correlation results for saliency maps, and to a lesser degree the last few layers, was revealed by the elementwise encoding method. While saliency maps were moderately predicted by low-, mid-, and high-level feature layers, the reverse did not hold. This shows that saliency integrates different feature activations but is rather agnostic to individual feature dimensions, which is in line with how saliency was previously defined. The same similarity analysis was performed with layers from the image classification and autoencoder networks and can be seen in Figures F.14A-D.

To visualize and quantify the notion of how different models and layers form clusters, we applied two techniques: Multidimensional Scaling (MDS) [76, 77] and Agglomerative Clustering. The former embeds pairwise similarities in a two-dimensional space and can thus be intuitively depicted. The latter is a hierarchical clustering algorithm that recursively merges the most similar data points until a desired number of clusters is reached. Here, the maximum distance between all elements in a cluster, starting from one data point per cluster, was used as a linkage criterion. Both techniques confirmed that the saliency model outputs form two separate clusters with larger distances between elements of the low-level compared to the high-level cluster (see Figures 3E and 3G). Furthermore, similarities between neighboring layers of MSI were highest and indicate a transformation towards a saliency map that is distinct from individual encoder features (see Figures 3F and 3H). The clustering analysis thus provided quantitative evidence for the observations made before. It also allowed us to reduce the computational costs of further analysis steps by selecting a subset of eight layers without sacrificing the large breadth of unique representations inherent to a single model. Again, the same cluster analysis was performed with layers from the image classification and autoencoder networks and can be seen in Figures F.14E-H, along with a description of the results in Appendix F.

### 3.2. Low- vs. High-Level Saliency Models

In light of the methodological differences and quantitative dissimilarities between low- and high-level saliency models, we contrasted their predictive accuracy with respect to fMRI data. To that end, we established a direct link between the eight saliency models and neural betas by means of three methods: RSA, ENC, and PRF (see Sections 2.7 and 2.8 for details). They either relate dissimilarity matrices of conditions for model and brain (RSA) or predicted and measured activations per voxel after fitting regression weights from entire saliency maps (ENC) or localized receptive fields (PRF). An agreement between results from these different but complementary approaches would thus render the conclusions more robust than findings from a single method.

We first applied all methods to the saliency maps from each of the eight models separately. This resulted in one Pearson correlation score per subject and ROI, which denotes the representational similarity across the entire neural population (RSA) or the median across individual voxels (ENC, PRF). We repeated this procedure for all five ROIs and averaged the scores across eight subjects (see Figure 4A). Notably, all saliency models and methods achieved a significant mean correlation (see Section 2.9 for details) with the corresponding neural targets. Performance scores were either normalized by the lower bound of the leave-one-subject-out ceiling (RSA) or the voxelwise noise estimates (ENC, PRF). This transformed values into the range between 0 and 1 and therefore enabled a fair comparison across methods and ROIs. What can be seen from the results is that individual high-level models consistently outperformed low-level ones in all ROIs except for early visual cortices. There, AIM achieved a mean correlation at the same level (RSA) or higher (ENC, PRF) than the best deep learning models and ICF surpassed SAM on one of the three methods (RSA). The dominance of AIM in the early ROI might be explained by the sparse and independent feature basis that was extracted from natural images via *independent component analysis* (ICA) and gave rise to saliency map predictions. Such features were suggested to resemble receptive field properties of simple cells in this area [78]. Among the high-level approaches, MSI attained the closest fits with neural data and was hence used as an exemplary model during later analysis steps. Notably, the same model performed best on the saliency benchmark (see Table B.2), which implies that the task of saliency prediction might align with neural computations. An overview of significance tests for pairwise model comparisons from Figure 4A can be seen in Figure G.15.

**Figure 4:**
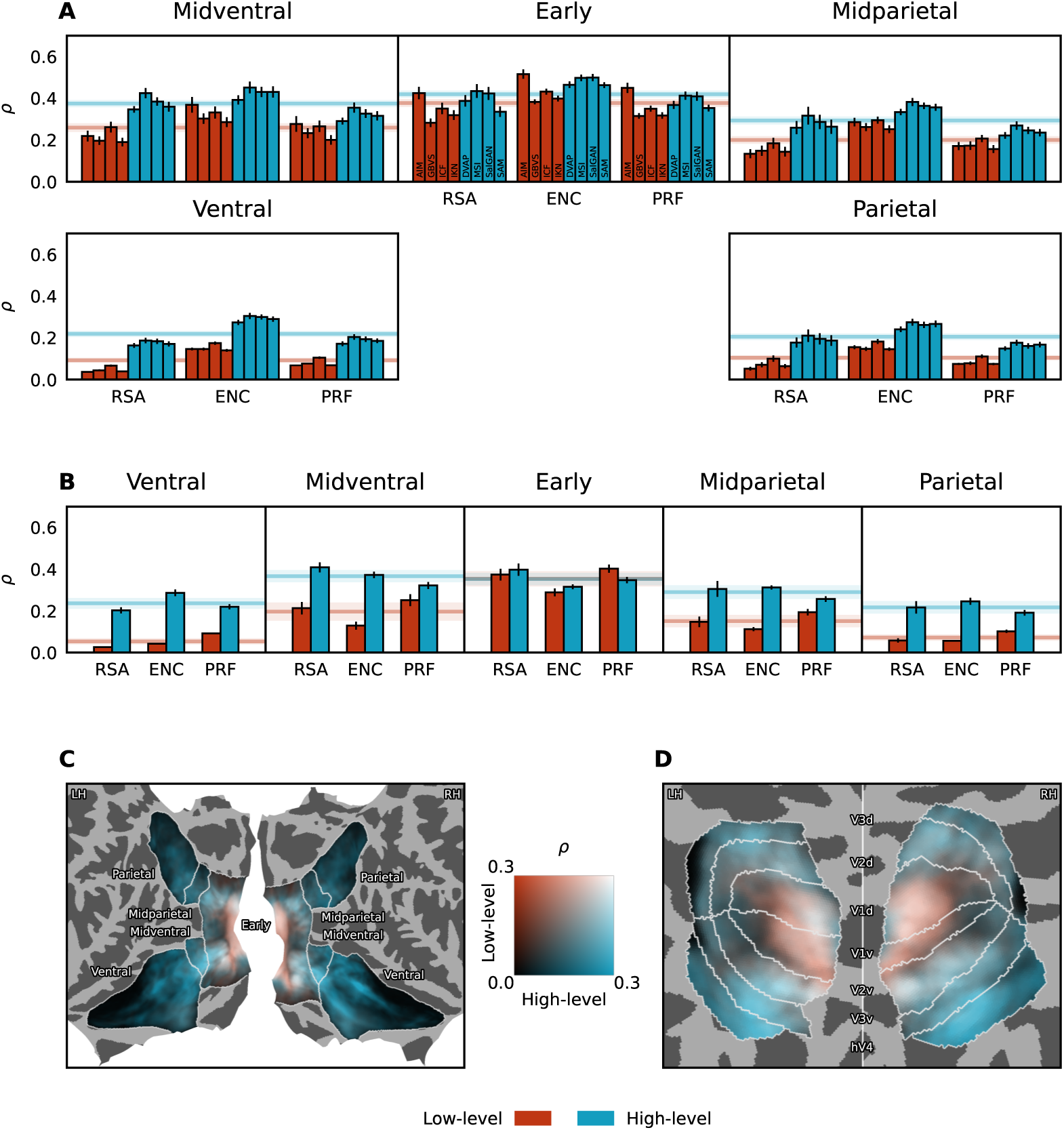
Group-average comparison between low- and high-level saliency models. (**A**) Mean correlation performance for each model, method, and ROI after noise normalization. The vertical line at the tip of a bar indicates the standard error. Model names associated with each bar are depicted in the top middle panel. Shaded horizontal lines represent the correlation coefficient after averaging the individual results per ROI across subjects, methods, and models within either the low- or high-level category. The shaded area around this mean represents the two-tailed 95% confidence interval. All individual model performances are significant. (**B**) Results based on the two grouped and disentangled model categories. Correlation scores ρ denote the relative contribution of low- and high-level saliency models to the predictive performance after fitting a joint encoding model via banded ridge regression. All grouped model performances are significant, as are the pairwise comparisons between categories for each ROI and method. (**C**, **D**) Group-average correlation scores per vertex on fsaverage. The two-dimensional colormap depicts the unique contribution of each feature space to the jointly estimated model. Disentangled correlation scores ρ were first obtained per subject, projected to a common brain space, and then averaged. Afterwards, the two visual streams (**C**) and retinotopically defined areas (**D**) were visualized on a flatmap and sphere respectively. Note that scores were not normalized by the noise ceiling here.

Overall, the mean correlation score per model category, summarized as the average over low- or high-level models across methods and indicated by the shaded horizontal lines in Figure 4A, was significantly highest in the early ROI and significantly decreased along the processing hierarchy of either visual stream. The progression along the ventral stream appeared to be mirrored in the dorsal stream, although midventral activations, which correspond to hV4, were significantly better predicted than midparietal ones. The three methods were in remarkably close agreement with respect to the evaluation of model performance and thus rendered the observations fairly robust. In general, the correlations of high-level models with fMRI data in early and midventral ROIs were surprisingly high given the simplistic representation of saliency maps. This could hint towards a meaningful and neurally relevant role of saliency. The difference between the average low- and high-level scores across methods was slightly but significantly more pronounced in the ventral compared to the parietal stream, and lowest in the early visual cortex. It must further be noted that the features giving rise to saliency maps were not independent between the two model categories and therefore contained spurious correlations.

For that reason, we then quantified the unique contribution of low- and high-level models to the predictive accuracy. First, the saliency maps from all four models within a category were grouped (see Sections 2.7 and 2.8 for details). Second, the two feature spaces were disentangled by means of a whitening procedure (RSA) or fitting a joint model using banded ridge regression (ENC, PRF). The results of this analysis can be seen in Figure 4B. Both model groups demonstrated a significant contribution to the mean correlation score for each method and ROI. Similar to before, the high-level feature space significantly exceeded the results from the low-level feature space starting from mid-level regions along the two visual streams. The difference in relative contribution was also significantly larger in (mid)ventral compared to (mid)parietal ROIs. Taken together, the high-level group contributed most to explaining neural data in the midventral area. This suggests that hV4 represents factors, such as a conjunction of complex visual features, that could account for the distinction of simplistic from naturalistic eye movement patterns. In early visual cortices, the mean correlation result was significantly equal between categories, albeit at a high performance level. Since the early ROI comprises areas from V1 to V3, it could hence benefit from a more fine-grained analysis of the contrast between low- and high-level models.

To that end, we plotted the voxelwise predictions from the banded ridge regression encoding model on fsaverage for all subjects (see Figure 4C and Figure 4D). The disentangled correlation scores revealed a clear distinction between V1, where activity was significantly better predicted by low-compared to high-level models, and V2/V3, where the opposite was true. The midventral area hV4 appeared to be particularly well predicted and dominated by the high-level saliency maps. This dominance continued along the lingual and fusiform gyri within the ventral region. High-level saliency maps also achieved significantly superior performance in all remaining areas of both visual streams but prediction scores generally degraded for either model category towards late processing stages. Correlation results along the inferior temporal gyrus were especially poor according to our analysis. This is likely due to a low signal-to-noise ratio evident from the noise ceiling estimates in those areas (see Figure I.17).

Low-level saliency hence only played an important role in area V1. However, neural activity within V1 for receptive fields located at eccentricities up to 0.5 visual degrees (see Figure K.19) was better predicted by high-level saliency models, although not significantly. This might be due to objects being preferentially located at the center of the screen (see Figure E.13), where subjects were instructed to fixate at all times throughout the experiments of NSD. Object-based saliency would thus be most prevalent at low eccentricities. This gave rise to a gradient in V1, where the major contribution to the joint prediction model switched from the high- to the low-level space at increasing eccentricities. To rule out that an image-independent center bias across the high-level saliency maps could explain this observation, we removed any spatial prior from the predictors (see Section 2.6). These results might therefore indicate an influence of true semantic saliency on even the earliest regions of the visual cortex.

### 3.3. Task Comparison of Deep Neural Networks

Beyond the comparison with low-level approaches, high-level saliency models can be characterized in relation to other deep learning networks. We selected MSI as the reference saliency model due to its superior performance with respect to both neural and saliency predictions over the other models of the high-level category. Furthermore, its backbone is based on the architecture of VGG16 with weights initialized from the original image classification task. This naturally allowed us to compare the influence of a new training task on the neural plausibility of emergent representations throughout the entire network. We thus not only considered the final output of a model but instead concatenated eight layers across its representational space (see Section 3.1 for details on the selection procedure). Neural predictions were then generated by fitting a separate voxelwise encoding model for MSI and VGG. The difference of correlation scores with neural targets indicated which cortical regions were best explained by each task and therefore revealed a functional map of visual areas.

Figures 5A and 5B demonstrate a comparison between MSI and VGG for voxels that were significantly predicted by at least one of the two networks. Differences in correlation scores were generally small, which suggests that features tuned towards saliency prediction did not drastically change with respect to the initial image classification task. Nevertheless, the percentage of voxels that correlated higher with one over the other network clearly differed across ROIs. VGG achieved better results in both the ventral and parietal ROI, where semantic information might be particularly relevant. In the midventral area, MSI compared favorably to the image classification network and thus corroborated the potential role of hV4 for saliency computations. This model advantage also held in early visual cortex and the midparietal ROI, albeit to a lesser degree. All model comparisons, expressed as the percentage of voxels assigned to either of the two models, are significant (see Section 2.9 for details).

**Figure 5:**
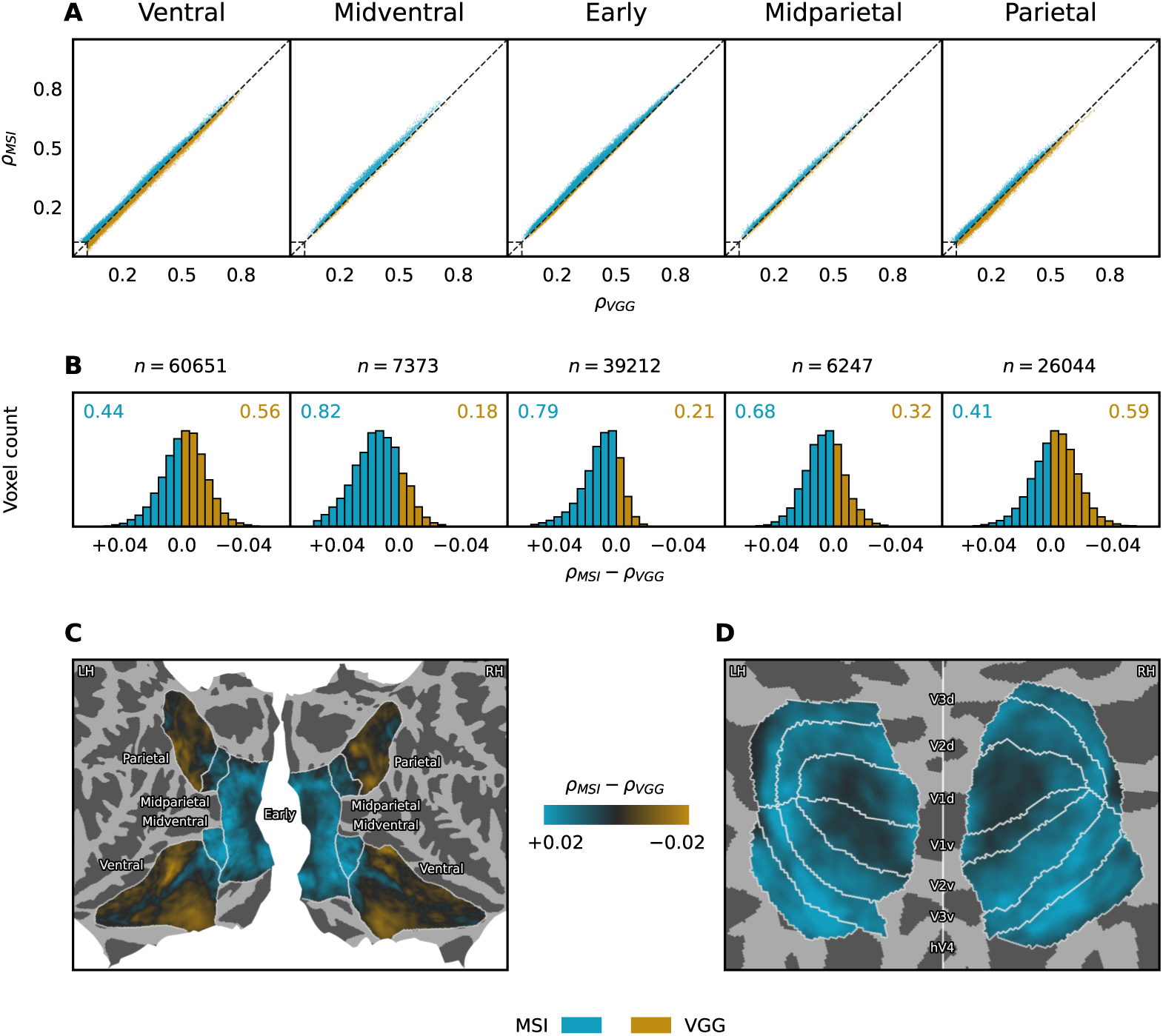
Model comparison between MSI and VGG with respect to the difference in correlation scores after fitting the two networks independently via ridge regression. (**A**) Voxelwise correlation results for the two models across subjects and per ROI, where points above or below the diagonal indicate an advantage for MSI or VGG respectively. The rectangles in the bottom left corner of the plots depict the minimum correlation score required for significant results. (**B**) A histogram that aggregates the correlation score differences between the models across all subjects and voxels, with *n* denoting the total number of significant voxels per ROI. The colored numbers at the top corners of the plots summarize the ratio of voxels that were best predicted by each model. All model differences are significant. (**C**, **D**) Group-average correlation differences between MSI and VGG per vertex on fsaverage. The colormap depicts the magnitude of a model advantage. Voxelwise correlation scores ρ were first obtained per model and subject, projected to a common brain space, and then averaged. Afterwards, the two visual streams (**C**) and retinotopically defined areas (**D**) were visualized on a flatmap and sphere respectively. Note that scores were not normalized by the noise ceiling here.

A fine-grained visualization of the group-average encoding results projected to fsaverage can be seen in Figures 5C and 5D. In addition to the previous observations, a more nuanced pattern within the early, ventral, and parietal areas became evident. First, the difference of correlation scores between the models was significantly less pronounced in V1 compared to V2/V3. Second, a considerable number of vertices along the inferior temporal lobe was better predicted by MSI in both hemispheres. This comprises the fusiform gyrus and posterior parts of the lingual gyrus. Although ventral regions have commonly been characterized by transformations akin to image classification networks [26], a subset of neural activity appeared to be more related to saliency-specific computations. Finally, a cluster of vertices at a late stage of the dorsal processing stream, roughly corresponding to the areas IPS3, IPS4, and IPS5 (localized via the atlas by Wang et al. [79]), displayed similar results. This indicates that a neural representation of saliency may also be reflected in parietal regions, which could be linked to oculomotor decision processes through interactions with FEF [80].

It must be noted that MSI and VGG not only differ by training task but also their architecture and number of learned weights associated with the selected layers. To rule out that the encoder-decoder structure of MSI alone can explain the observed differences in predictive performance, we further contrasted the scores with an autoencoder (see Figure H.16). Overall, the correlation differences were considerably higher than before and almost all voxels or vertices were assigned to the saliency network. This suggests that the autoencoder may not be a very suitable candidate model of cortical visual processing. In V1, the differences between the two networks were relatively small, which hints towards the notion that image reconstruction relies on low-rather than high-level features.

### 3.4. Layer Mapping of Deep Neural Networks

Next, we wanted to narrow in on the model comparison between MSI and VGG by regarding the assignment of individual network layers to voxels. This allows for a finer mapping of model representations to cortical activations and can generate additional insights about the intermediate processing stages of saliency computations. One problem is that spurious correlations between layers impede the analysis of their unique contribution to the prediction of neural activity [81]. We hence treated layers as individual feature spaces and applied banded ridge regression towards a disentangled evaluation of layer preferences per voxel. To limit the computational costs of this approach, we reduced the number of feature spaces by selecting a subset of layers per model that was deemed particularly relevant. More specifically, a joint encoding model based on eight layers of a given network (see Section 3.1 for details on the selection procedure) was estimated and four layers with the highest individual contributions across ROIs were kept for further analysis steps. The decomposed correlation scores of each feature space served as the basis for a winner-take-all (WTA) mechanism, where one layer was assigned per voxel. This procedure was performed for MSI and VGG separately (see Figure 6A and 6B). The ratio of significant voxels that were best explained by a layer within one ROI and across subjects indicated its importance towards accurate prediction results.

**Figure 6:**
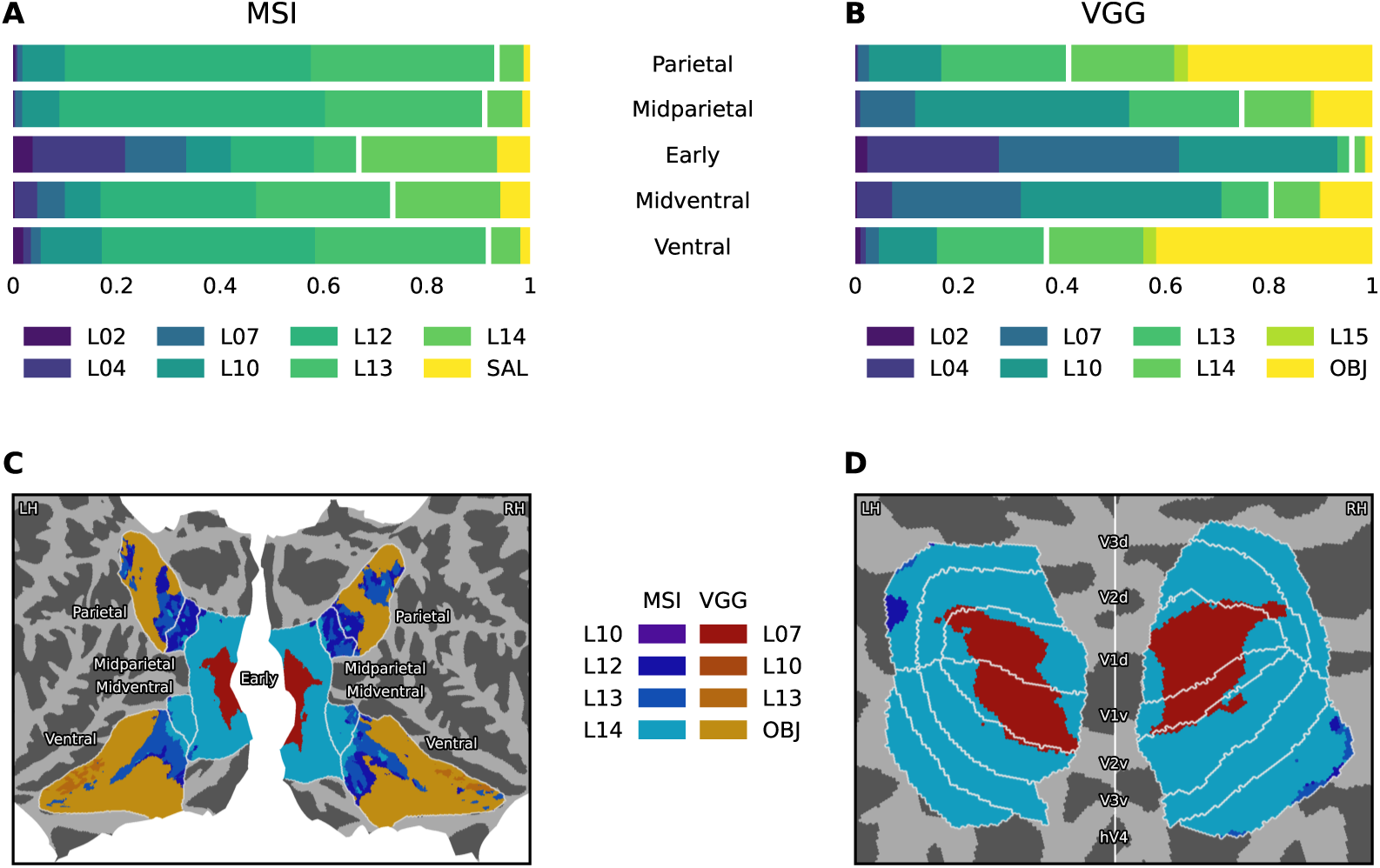
Quantification of layer preferences for the selection of representative network layers and the visualization of finer-grained model differences. (**A**, **B**) Group-average ratio of voxels that were best predicted by each layer after fitting a joint encoding model per network, subject, and ROI. The disentangled evaluation of individual predictors from either MSI (**A**) or VGG (**B**) was limited to the previous selection of eight layers. Associated colors in the horizontal bars are arranged from left to right regarding their position within the network hierarchy. White gaps indicate the split between encoder and decoder layers (**A**) or convolutional and dense layers (**B**) occurring after L13. Note that all layers in MSI up to and including L13 were initialized from VGG weights and fine-tuned on a saliency prediction task. Subsequent layers are more task-specific and culminate in the model output of saliency maps (SAL layer) in MSI or object labels (OBJ layer) in VGG respectively. Four layers with the highest individual contributions across ROIs were selected per model. (**C**, **D**) Group-average layer preference per vertex on fsaverage. The colormap depicts which of the eight layers from the extracted subset of either MSI or VGG was assigned based on predictions by a jointly estimated model. Disentangled correlation scores ρ were first obtained per subject, projected to a common brain space, averaged, and then evaluated with respect to model and layer preferences. Afterwards, the two visual streams (**C**) and retinotopically defined areas (**D**) were visualized on a flatmap and sphere respectively. Note that scores used for the layer mapping were not normalized by the noise ceiling here.

In the case of MSI, saliency maps did not substantially contribute to the joint encoding model performance in any of the tested ROIs but achieved their best results in the early and midventral area. Likewise, the penultimate layer L14 was particularly relevant in the same ROIs, albeit with a much stronger contribution among the predictors. Note that the two layers L14 and SAL represent the decoder of MSI and thus more task-specific features, which explained neural activity in early and midventral ROIs reasonably well. The early visual cortex was also characterized by early layers of MSI, whereas intermediate network layers contributed substantially to the prediction of neural beta values in all other areas. Overall, layers L10, L12, L13, and L14 of MSI were most relevant across ROIs and kept as a subset of predictors for model comparison. The same selection procedure was conducted for VGG, where layers L07, L10, L13, and OBJ dominated the predictions of a joint encoding model. The VGG analysis revealed a gradient of representations along the two visual streams. Early feature maps of VGG corresponded to early visual cortices, while object representations from the final stages of the network hierarchy were associated with ventral and parietal areas. Altogether, the fine-tuning of features originating from VGG towards saliency prediction in MSI rendered intermediate network layers more important. This came at the cost of earlier layers and supports the notion that complex visual representations with a topographic organization, akin to high-level convolutional feature maps, may be particularly relevant for saliency computations in the brain.

To conduct a more fine-grained comparison between MSI and VGG than in Section 3.3, we quantified the relative contribution of layers across the two networks via banded ridge regression. Feature spaces consisted of the four most representative layers from each model and were jointly fitted on the neural data. Although the two selected sets of layers are not directly comparable nor specific to an ROI, they still allow us to localize regions where features with an overall strong predictive performance are uniquely relevant. Figures 6C and 6D demonstrate the assignment of individual layers to vertices that were significantly predicted by at least one of the eight layers. For evaluation, we first summed the decomposed correlation scores across the four layers from each network, determined the winner, and then chose a layer from the best model with the highest individual performance via the WTA technique. This procedure was intended to characterize both the model and layer preferences of individual vertices. The topmost layer from MSI under consideration (L14) was assigned to the majority of vertices in V2, V3, and hV4, whereas V1 primarily corresponded to representations of L07 from VGG. Notably, parts of V1 with receptive fields located at eccentricities up to 0.5 visual degrees (see Figure K.19), were better predicted by the high-level saliency model, similar to the results in Figures 4D and 5D. We also observed that object representations of layer OBJ from VGG dominated most regions of the ventral and parietal ROIs. Nevertheless, intermediate layers from MSI were assigned to vertices along the fusiform gyrus and posterior parts of the lingual gyrus. This model advantage was also found in the midparietal ROI and areas within IPS0, IPS1, and IPS2 in addition to IPS3, IPS4, and IPS5 of the parietal lobe (localized via the atlas by Wang et al. [79]). Generally, the locations along both visual streams were in close agreement with the results from Figure 5C.

Figures J.18A and J.18B further depict the magnitude of model preferences obtained from the disentangled evaluation procedure. The distribution and relative strength of model advantages over cortical regions was approximately equal to the whole-network analysis in Figure 5C, with the exception of V1. There, VGG achieved a higher aggregate performance among its four layers in comparison with MSI. This illustrates that our selection of layers was largely representative of the corresponding models. To shed more light on the layer profile of individual vertices, we visualized the assignment of second-best layers in Figures J.18C and J.18D and described the results in Appendix J.

### 3.5. Topographic Representation of Saliency Maps

Finally, we investigated whether saliency maps might indeed be explicitly represented and topographically organized in the brain. Our underlying assumption is that if a single saliency map exists as suggested by computational models, it should be localizable via pRF mapping. This is a reasonable requirement given that saliency maps must be tightly linked to oculomotor control and thus preserve the spatial properties of the visual field. Here, we first estimated pRF locations based on the saliency maps generated by MSI (see Figure 7). Polar angle estimates in V1, V2, V3, and hV4 aligned well to the results obtained from NSD (see Figure K.19 for comparison) and accurately traced the borders of these areas. Interestingly, vertices in the ventral and parietal surface ROIs of both hemispheres were found to have receptive field centers preferentially located in the upper and left quadrants of the visual field (see Figure 8A). This is in stark contrast to the contralateral visual field preference in the left and right hemisphere for parietal regions, and to a lesser degree ventral regions, as evident in NSD. Saliency or, more generally, visuospatial attention might hence be subjected to a bihemispheric spatial bias in these areas unlike general neural processing of visual stimuli. To test whether this attentional bias could have an effect on downstream tasks, we analyzed the polar angle distribution of the first fixation in comparison to all remaining fixations for two datasets independent from NSD (see Figure 8B). Both revealed a preference for the upper and left quadrants of the visual field as well but only for the first fixated location. This suggests that the spatial bias of receptive field locations related to saliency in the ventral and parietal ROIs may be linked to oculomotor control.

**Figure 7:**
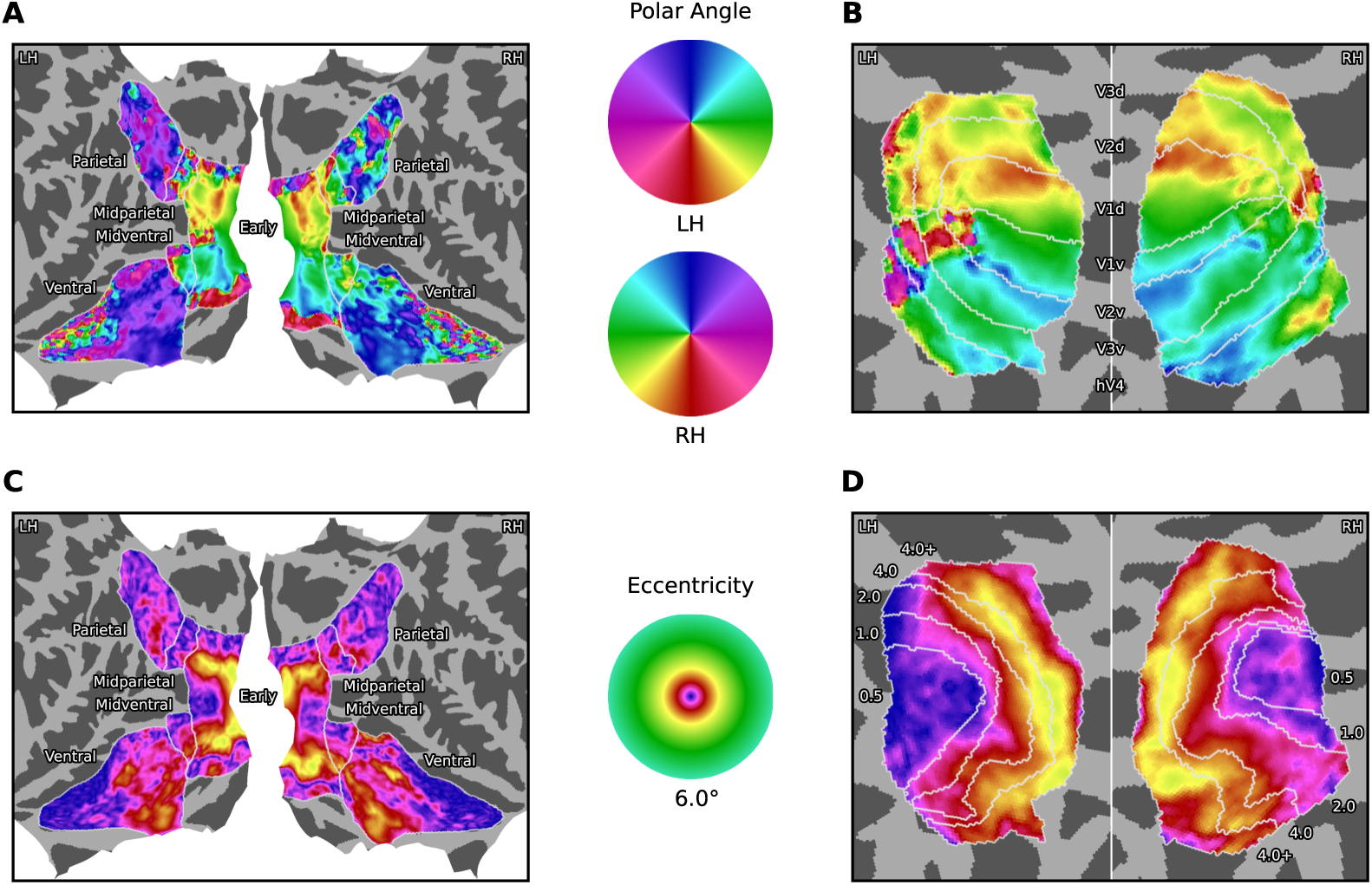
Group-average polar angle and eccentricity estimates from pRF mapping based on saliency maps of MSI per vertex on fsaverage. Voxelwise receptive field locations were first obtained per subject, projected to a common brain space, averaged, and then converted to polar angle (**A**, **B**) and eccentricity (**C**, **D**) values. Afterwards, the two visual streams (**A**, **C**) and retinotopically defined areas (**B**, **D**) were visualized on a flatmap and sphere respectively. Borders on the sphere visualization for eccentricity estimates (**D**) are based on the eccentricity delineation provided by NSD. Note that any spatial bias was removed from the predictors prior to pRF mapping.

**Figure 8:**
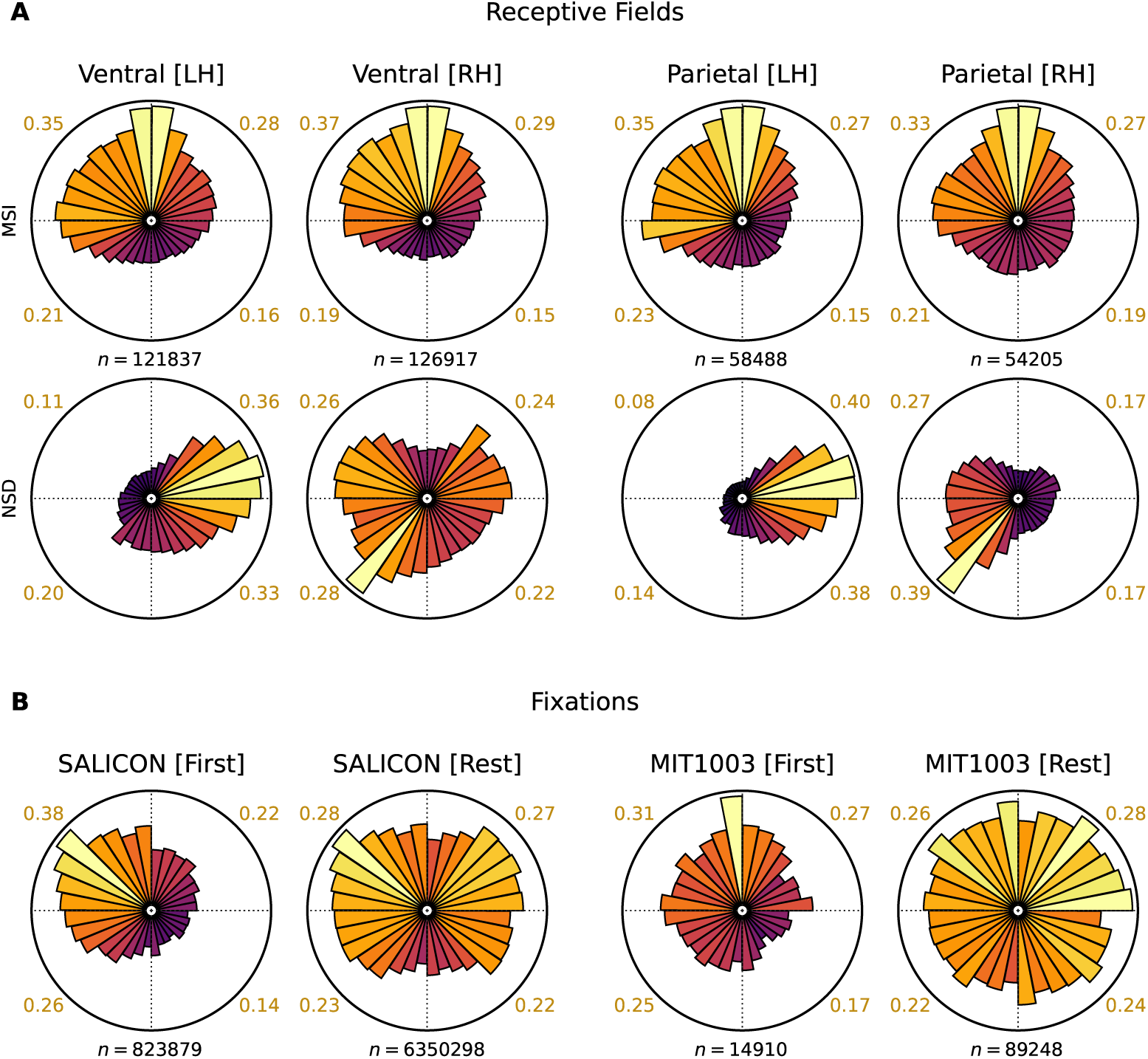
Polar angle histograms of estimated receptive field locations (**A**) or empirical fixation locations (**B**). Panels depict the visual field divided into 32 angular bins. The radius of each wedge is proportional to the number of elements within the corresponding polar angle interval at any eccentricity. Dotted lines divide the visual field into four quadrants and colored numbers outside the plots represent the percentage of data points that are located in the respective quadrant. (**A**) For each ROI and hemisphere, *n* equals the total number of significant vertices across the receptive field estimates of individual subjects. The two rows contrast our results from pRF mapping of MSI saliency maps to neural data with the estimates provided by NSD. Note that any spatial bias was removed from the predictors prior to pRF mapping. (**B**) Two datasets with empirical data were analyzed: SALICON [44] and MIT1003 [82]. The former recorded mouse movements as an accurate proxy for gaze information and the latter is based on eye-tracking data. Both were acquired under a free-viewing paradigm starting from a central position within natural images. Fixation measurements were divided into the first and all remaining instances. For each plot, *n* equals the total number of fixations that were aggregated across stimuli and subjects.

Furthermore, we observed smaller eccentricity values throughout all ROIs for pRF mapping based on MSI saliency maps compared to the NSD estimates. The increased number of saliency-selective receptive fields dedicated to central parts of the visual field might explain previous results, where prediction accuracies from high-level saliency compared favorably to several contrasting approaches at low eccentricities within early visual areas (see Figures 4D, 5D, and 6D). Figure L.20 depicts receptive field sizes and fits obtained from the same mapping procedure. Receptive field sizes, as defined by the standard deviation of the best-fitting Gaussian, increased along the two streams even with respect to saliency maps. Correlation scores at low eccentricities in early visual regions were comparatively small but otherwise demonstrated an accurate prediction of neural activity in V1, V2, V3, and hV4. Notably, high correlation results within gyri of the temporal lobe corresponded to the same areas that were previously suggested to partake in saliency computations.

The question still remains whether these results were due to general visual features that have shown to give rise to topographic mappings [67] or salient information that is independent from individual feature channels. For that reason, we estimated pRF parameters using banded ridge regression, where the eight selected layers of MSI each formed a separate feature space. A subset of 1,000 unique images was chosen per subject to reduce the computational load of this approach. The disentangled polar angle and eccentricity estimates per layer can be seen in Figure 9 and Figure 10 respectively. Although all layers preceding the final output of MSI gave rise to a topographic mapping in the early and midventral ROIs, saliency maps themselves failed to do so. Especially layers L12, L13, and L14 yielded results that were in good agreement with pRF estimates from NSD. This suggests that saliency maps may not be exclusively represented in these regions but rather arise from localized feature computations. A remarkable observation was that the aforementioned spatial bias towards the upper and left visual field only became evident for saliency maps and not any of the preceding layers. This phenomenon might therefore be specific to visuospatial attention and possibly linked to the fixation biases in Figure 8B. Receptive field sizes varied for intermediate layers (see Figure M.21) but were somewhat constant for saliency maps throughout all cortices. Regarding the correlation scores of individual feature spaces, early model layers achieved high fits in the early ROI, whereas intermediate and late model layers further captured the neural activity in parts of the ventral and parietal stream (see Figure N.22). The unique contribution of saliency maps was generally small but, once again, remarkably pronounced in bihemispheric gyri along the temporal lobe. Overall, the juxtaposition between the topographic mapping results of saliency maps with and without disentanglement showed that early and midventral areas might primarily be organized according to general visual features instead of saliency. This would assign saliency computations in these regions a secondary but yet important role.

**Figure 9:**
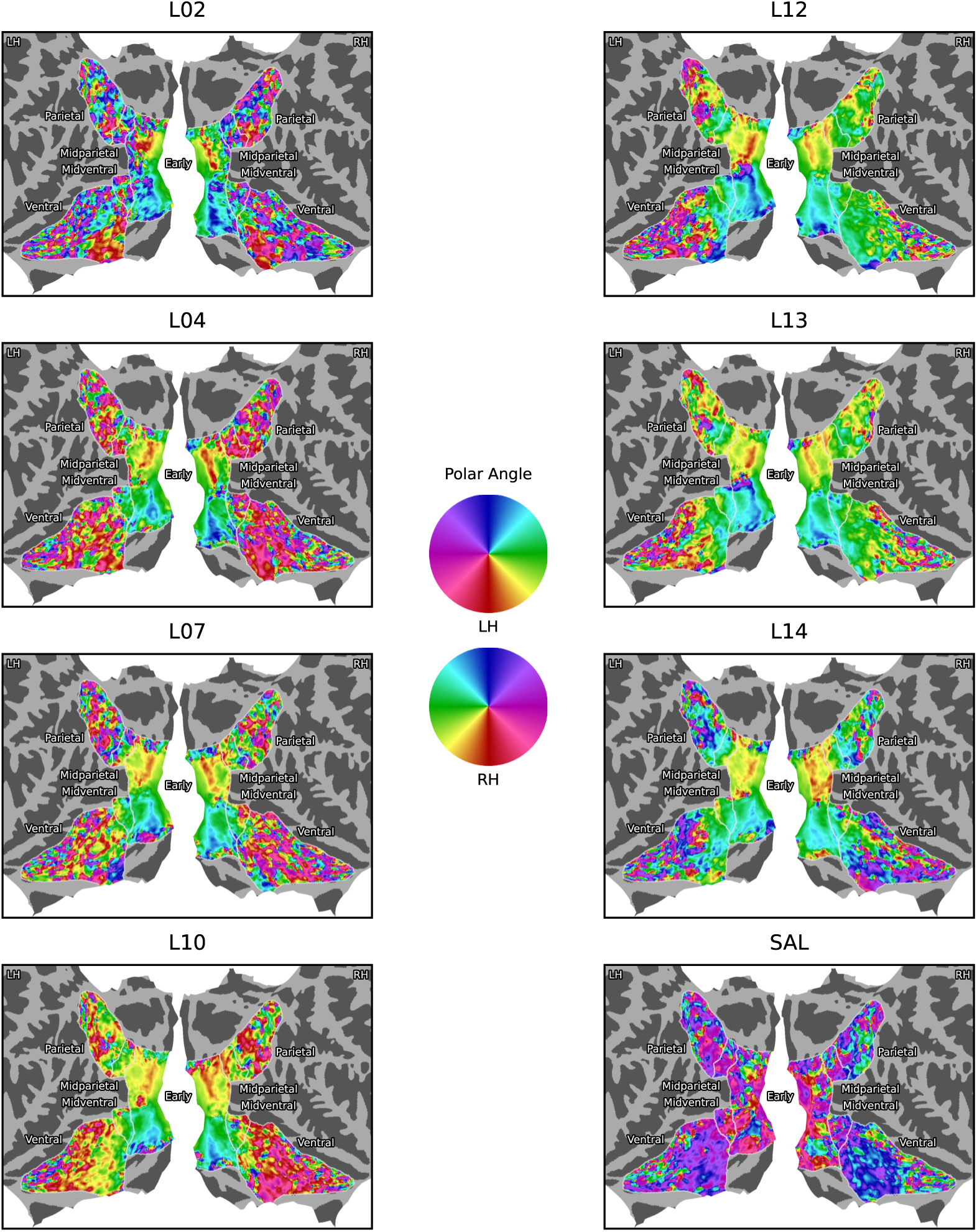
Group-average polar angle estimates from pRF mapping based on a disentangled subset of eight MSI layers per vertex on fsaverage. Voxelwise receptive field locations were first obtained per subject, projected to a common brain space, averaged, and then converted to polar angle values. Afterwards, the two visual streams were visualized on a flatmap for each layer separately. Note that any spatial bias was removed from the predictors prior to pRF mapping.

**Figure 10:**
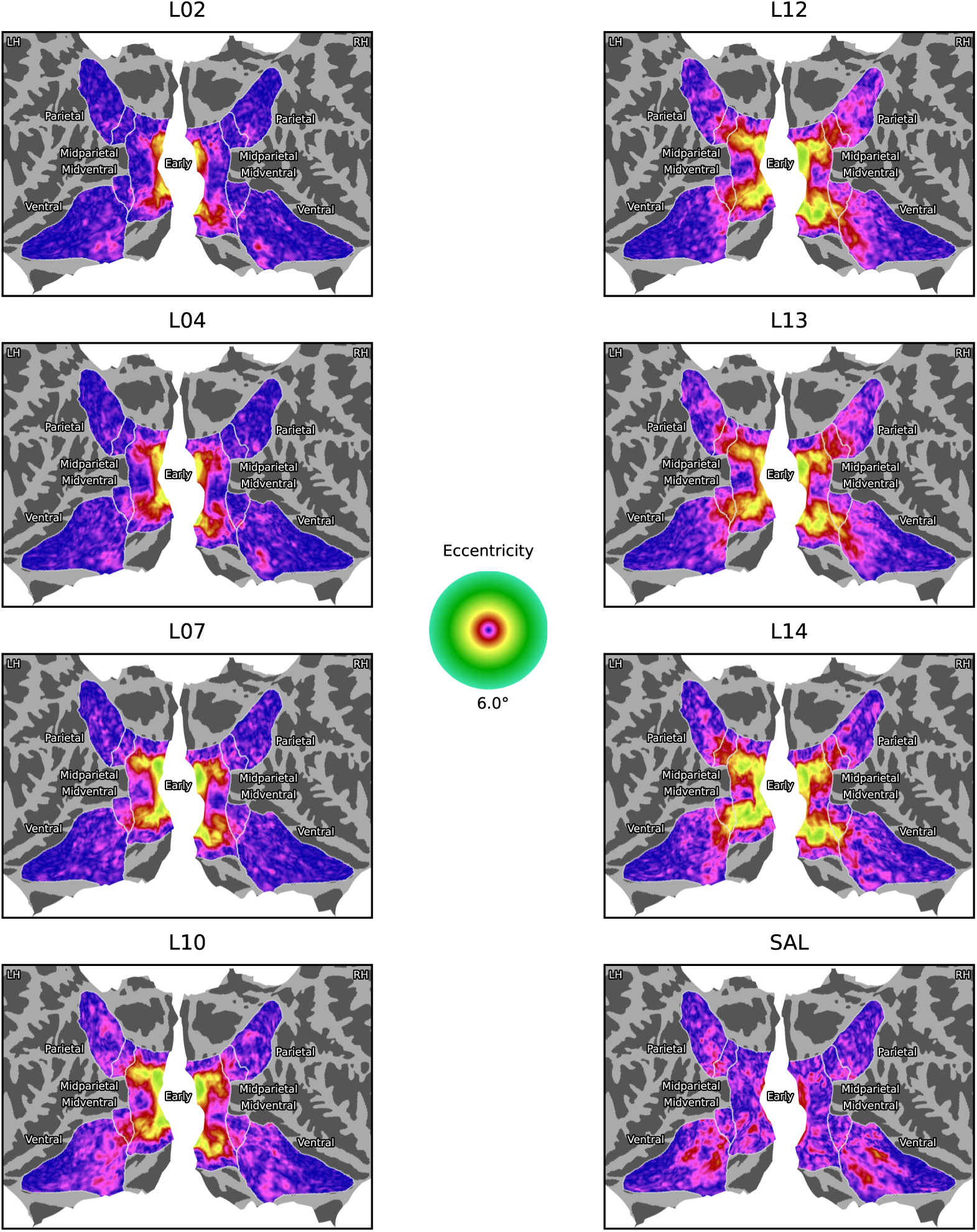
Group-average eccentricity estimates from pRF mapping based on a disentangled subset of eight MSI layers per vertex on fsaverage. Voxelwise receptive field locations were first obtained per subject, projected to a common brain space, averaged, and then converted to eccentricity values. Afterwards, the two visual streams were visualized on a flatmap for each layer separately. Note that any spatial bias was removed from the predictors prior to pRF mapping.

## 4. Discussion

Over the course of this study, we investigated the neural correlates of high-level saliency by contrasting it with low-level approaches, neural networks trained on different tasks, and general visual features at multiple levels of abstraction. The techniques that were utilized to link model representations to the brain consisted of RSA, a voxelwise encoding model, and pRF mapping that all either disentangled feature spaces or regarded them separately. Our analyses were aimed at characterizing the cortical representation of saliency in response to naturalistic stimuli from NSD and yielded remarkable agreement across methods. The results suggest that hV4, which corresponds to the midventral ROI, plays a central role for the processing of information that encodes saliency of complex visual scenes. Later stages along both the ventral and dorsal stream hierarchy were critically involved as well. Overall, saliency models with superior performance on a computer vision benchmark were also more accurate predictors of cortical representations. High-level saliency computations thus appear to constitute an ecologically valid task of the human visual system.

These observations resulted from a series of research questions revolving around the cortical representation of high-level saliency. Our first question addressed the differences between low- and high-level saliency and revealed that all ROIs beyond V1 were significantly better predicted by high-level models. Their differential ability to predict neural activity was particularly evident in (mid)ventral areas (see Figure 4), which confirmed our hypothesis that ventral processes are necessarily involved through their spatial encoding of semantic information. Next, we studied the differences between a high-level saliency model and deep neural networks trained on other visual tasks. Our results showed that neural activity in hV4 appeared more akin to feature maps learned by a model that is tuned towards saliency prediction than image classification or reconstruction (see Figures 5 and H.16). We observed this model advantage in visual areas V2 and V3 as well. These findings support the notion that saliency computations, as a representation subserving eye movement plans, are neurally relevant. Indeed, the stimulus-driven guidance of eye movements towards interactions with our environment is a fundamental task of visual processing [83].

Summarizing these intermediate results, we clearly demonstrated the involvement of ventral visual processes with respect to saliency-specific computations although they have traditionally been linked to object recognition [26]. Some of the regions along the lingual and fusiform gyri were best explained by semantic saliency after disentangling its contribution from low-level models, neural networks trained on two different tasks, or intermediate layers of MSI (see Figures 4, 5, 6, and H.16). All remaining areas within the ventral ROI were best predicted by semantic features inherent to the final classification layer of VGG (see Figure 5). This suggests that some of the information processing carried out in the temporal lobe underlies saliency computations. The functional description of the fusiform and lingual gyri comprises face, body, and word recognition among other high-level visual processes [84]. These features in particular are attentionally relevant and have proven to reliably elicit saccadic eye movements [21]. This could distinguish their role from other category-selective activations that are better captured by the image classification network. Another possibility is that the topographic organization of high-level features in MSI gives rise to this observation. Unlike the image classification network, spatial information of semantic content is preserved throughout all layers in the saliency model.

Interestingly, ventral areas corresponded to object-specific layers in the saliency model, while midventral or early ROIs were aligned with more saliency-specific layers (see Figure 6). If the sequence of transformations in the model bears some resemblance with neural processing, then ventral feature extraction should precede and inform saliency computations in hV4. Complex visual features have indeed shown to guide fixation patterns in response to natural images. This requires object processing with an emphasis on face and text detection [21], typically associated with the temporal cortex. For that reason, hV4 may receive semantic information through anatomical backward connections from ventral areas [23] and combine them with earlier visual features into an abstract representation related to attentional selection [85]. This would establish a transformative hub in hV4 that may even incorporate top-down factors, such as task instructions [86]. To characterize the flow of information along the ventral processing stream, future studies could assign individual layers to voxels over time based on fMRI timeseries data rather than beta values.

Our final research question addressed whether cortical saliency is topographically organized and independent from specific visual features. Population receptive field mapping based on high-level saliency maps led to polar angle and eccentricity estimates in early and midventral regions as anticipated from prior measurements of visual field maps [87] and NSD experiments (see Figure 7). At first glance, these findings seemed to point towards an explicit representation of saliency maps in hV4. Mazer and Gallant [16] have also argued that this region contains a visually driven and topographically organized saliency map but considered it as part of a saliency network that comprises additional cortical and subcortical structures. While our results are in agreement with their findings and hint towards a key role of hV4 as well, its function appeared more complex and cannot be characterized independently of the features that give rise to a saliency map. A detailed analysis revealed that its neural profile resembled the penultimate layer of MSI, among a subset of eight layers, more closely than saliency maps themselves (see Figure 6). The disentangled pRF mapping estimates corroborated the notion that activity within this region is not sufficiently accounted for by saliency maps when controlling for visual features at multiple levels of abstraction (see Figures 9 and 10). Under these circumstances, a topographic organization could not be identified anymore and therefore we conclude that the function of hV4 cannot be characterized by the representation of a saliency map alone. It may instead compute and integrate salient features simultaneously and contribute to oculomotor planning processes elsewhere. As such, the cortical representations of saliency and general visual features are deeply entangled for naturalistic image viewing.

Besides these main conclusions derived from our three research questions, one striking observation in both the ventral and parietal ROI was made after estimating pRF parameters based on saliency maps. Receptive fields were preferentially located in the upper and left quadrants of the visual field within both hemispheres, whereas a contralateral bias could be seen in pRF runs of NSD using moving bar stimuli (see Figure 8). This phenomenon persisted even after saliency maps were disentangled from other visual features of MSI. A spatial bias towards the upper and left quadrants might therefore be specific to saliency. Leftward biases have also repeatedly been described in studies of spatial attention [88] and were commonly attributed to a dominance of right-hemispheric attention networks [89]. Neuroimaging data supported this notion and revealed a bihemispheric leftward bias as the result of enhanced attention-related information transfer in IPS from the right to the left hemisphere [90]. Our pRF mapping results based on saliency maps corroborate the finding that also the left hemisphere preferentially encodes left visual field attention in the parietal cortex. Interestingly, Dickinson and Intraub [91] have found a leftward bias even for first saccades under a free-viewing paradigm when subjects fixated centrally on images of natural scenes, which extends to a variety of experimental conditions [92]. We were able to confirm this finding on two datasets (see Figure 8), where the first fixation preferentially targeted the left and upper visual field while all remaining fixations were more evenly distributed. However, spatial cueing may reduce or even eliminate the attentional bias towards the left side [93]. Studies aimed at the topographic mapping of parietal cortices via attentionally demanding tasks observed a clear contralateral organization in both hemispheres instead [94, 95, 96]. This indicates that a leftward bias is rather related to naturalistic stimulus-driven saliency than top-down attentional processes and could exert an influence on fixation behavior.

Furthermore, our analyses suggested that a saliency-related bias towards the upper quadrants was even more pronounced. This was then confirmed in an eye-tracking dataset for the first saccade (see Figure 8). Kraft et al. [97] discovered the same vertical asymmetry in the frontoparietal attention network during a covert visual search task. In NSD, participants were instructed to perform continuous recognition aimed at memorizing previously shown images, while maintaining central fixation. One possible explanation is that they might have employed a viewing strategy reminiscent of covert visual search. While the precise reasons for the observed upward bias remain elusive, our results propose that an attentional preference for the upper left quadrant of the visual field would emerge under experimental conditions similar to NSD. The same applies to our findings regarding the reduced eccentricity of receptive fields throughout all cortices when mapping saliency maps. Unlike previous studies, we not only detected spatial biases of attention in posterior parietal areas but also the temporal lobe. This observation might, however, be limited to experiments involving naturalistic stimuli that require a more complex scene analysis through ventral visual processing. Our proposed link between the attentional bias and oculomotor control needs to be validated in future neuroimaging studies that include the first saccade after image viewing of natural scenes.

While saliency computations must incorporate semantic features, another requirement is that they inform oculomotor planning processes. Again, hV4 constitutes a likely candidate region by means of its robust projections to (mid)parietal areas [23]. This establishes an indirect connection from hV4 to the oculomotor system and would allow for saliency computations to guide eye movement behavior. Only in comparison with an image classification model did we observe the relevance of saliency in subregions of IPS but found no indication of explicit saliency computations in the dorsal pathway otherwise. Nevertheless, there is clear evidence that the interplay between IPS and FEF is crucial for attentional selection [98]. We thus argue that frontoparietal areas perform downstream computations based on salient features from ventral processing, similar to the account by Mazer and Gallant [16]. Our results did not reveal a topographic mapping of these areas with respect to saliency maps either. To detect a gradual representation of pRF angle and eccentricity estimates, a more attentionally demanding task may be required [94]. Another limitation is that receptive field sizes at late stages of the visual cortical hierarchy are large and therefore more difficult to map with pRF models when the field of view is small [68], as is the case in NSD. However, the rather symmetrical results between the ventral and parietal ROIs in our analyses suggest that they engage in cross-stream interactions.

Another unexpected finding was that high-level saliency best predicts activity in V1 at receptive field locations up to 0.5 visual degrees away from the center of fixation, whereas low-level saliency or intermediate VGG layers dominated at higher eccentricities (see Figures 4 and 6). The projections between areas along the ventral visual pathway have shown to primarily encode central parts of the visual field [99]. If we assume the existence of saliency computations in the ventral stream, then it should exhibit a bias towards foveal areas. Our observations thus suggest that semantic saliency, originating from subsequent stages of the ventral stream, may affect visual areas as early as V1. To conclusively test whether this interpretation of our results is indeed correct, future studies investigating the effective connectivity along the ventral visual pathway for both feedforward and feedback connections are needed. One limitation in our analyses is that only a subset of network layers was selected due to computational considerations. The assignment of layers to voxels and implications regarding the sequence of transformations in the brain should hence be regarded with caution.

In conclusion, our study implicates area hV4 as a key stage of saliency computations, linking low- and high-level features to attentional selection. We found that neural responses to naturalistic images were entangled with general visual features, which hints towards a transformative role rather than the sole representation of a saliency map independent from individual features. Besides hV4, we observed neural correlates of saliency computations distributed over areas along the dorsal and ventral pathway as well as early visual cortex. Our results suggest that concerted activity is required to encode where salient features are located in complex scenes. To that end, saliency maps generated by computational models encapsulate information that has shown to be neurally relevant and shed light on visual processes subserving ecologically valid tasks, such as eye movement control. Our results were obtained from a multitude of methods and contrasts applied to a large-scale fMRI dataset, which forms the first instance of linking deep neural networks trained towards accurate saliency prediction to the brain.

## Acknowledgement

This study has received funding from the European Union’s Horizon 2020 Framework Programme for Research and Innovation under the Specific Grant Agreement Nos. 785907 (Human Brain Project SGA2) and 945539 (Human Brain Project SGA3). We would like to thank Agustin Lage-Castellanos for the helpful discussions about banded ridge regression in combination with population receptive field mapping. Finally, we gratefully acknowledge the support of NVIDIA Corporation with the donation of a Titan X Pascal GPU used for this research.

## Appendix A. Streams ROIs

**Table A.1:** The delineation of visual streams based on the aggregation of ROI labels from the probabilistic atlas by Wang et al. [79] and manually drawn modifications by the authors of the NSD. This textual description was adapted from the ROI definitions provided in the NSD manual. Stream labels were originally defined on fsaverage and then transformed to subject-specific surface and volume versions.

| *ROIs* |  |
| --- | --- |
| Early | V1v, V1d, V2v, V2d, V3v, V3d |
| Midventral | inferior boundary of hV4 including the inferior occipital gyrus (IOG) and based on the anterior edge of the inferior occipital sulcus (IOS) |
| Ventral | anterior lingual sulcus (ALS) including the anterior lingual gyrus (ALG) and based on the inferior lip of the inferior temporal sulcus (ITS) as well as the midpoint of the occipital temporal sulcus (OTS) |
| Midparietal | V3A, V3B |
| Parietal | IPS0, IPS1, IPS2, IPS3, IPS4, IPS5, SPL1 |

## Appendix B. Saliency Model Benchmark

**Table B.2:** A ranking of model performances reported by the MIT/Tübingen saliency benchmark [40]. Low-level models (top) and high-level models (bottom) were evaluated on the MIT300 test set [100] using 6 common metrics. Arrows indicate whether higher or lower values are closer to the gold standard. Among all models, MSI achieved the highest results (bold) and high-level models consistently outperformed low-level models on all metrics but KLD.

|  | *AUC* ↑ | *sAUC* ↑ | *NSS* ↑ | *CC* ↑ | *KLD* ↓ | *SIM* ↑ |
| --- | --- | --- | --- | --- | --- | --- |
| AIM | 0.762 | 0.665 | 0.882 | 0.342 | 1.248 | 0.410 |
| GBVS | 0.806 | 0.630 | 1.246 | 0.479 | 0.888 | 0.484 |
| ICF | 0.833 | 0.696 | 1.613 | 0.588 | 0.708 | 0.558 |
| IKN | 0.781 | 0.632 | 1.113 | 0.430 | 0.961 | 0.465 |
| DVAP | 0.843 | 0.726 | 1.931 | 0.663 | 0.629 | 0.585 |
| MSI | **0.874** | **0.779** | **2.305** | **0.779** | **0.423** | **0.670** |
| SalGAN | 0.850 | 0.735 | 1.862 | 0.674 | 0.757 | 0.593 |
| SAM | 0.847 | 0.731 | 1.955 | 0.663 | 1.275 | 0.599 |

## Appendix C. Saliency Model Predictions

**Figure C.11:**
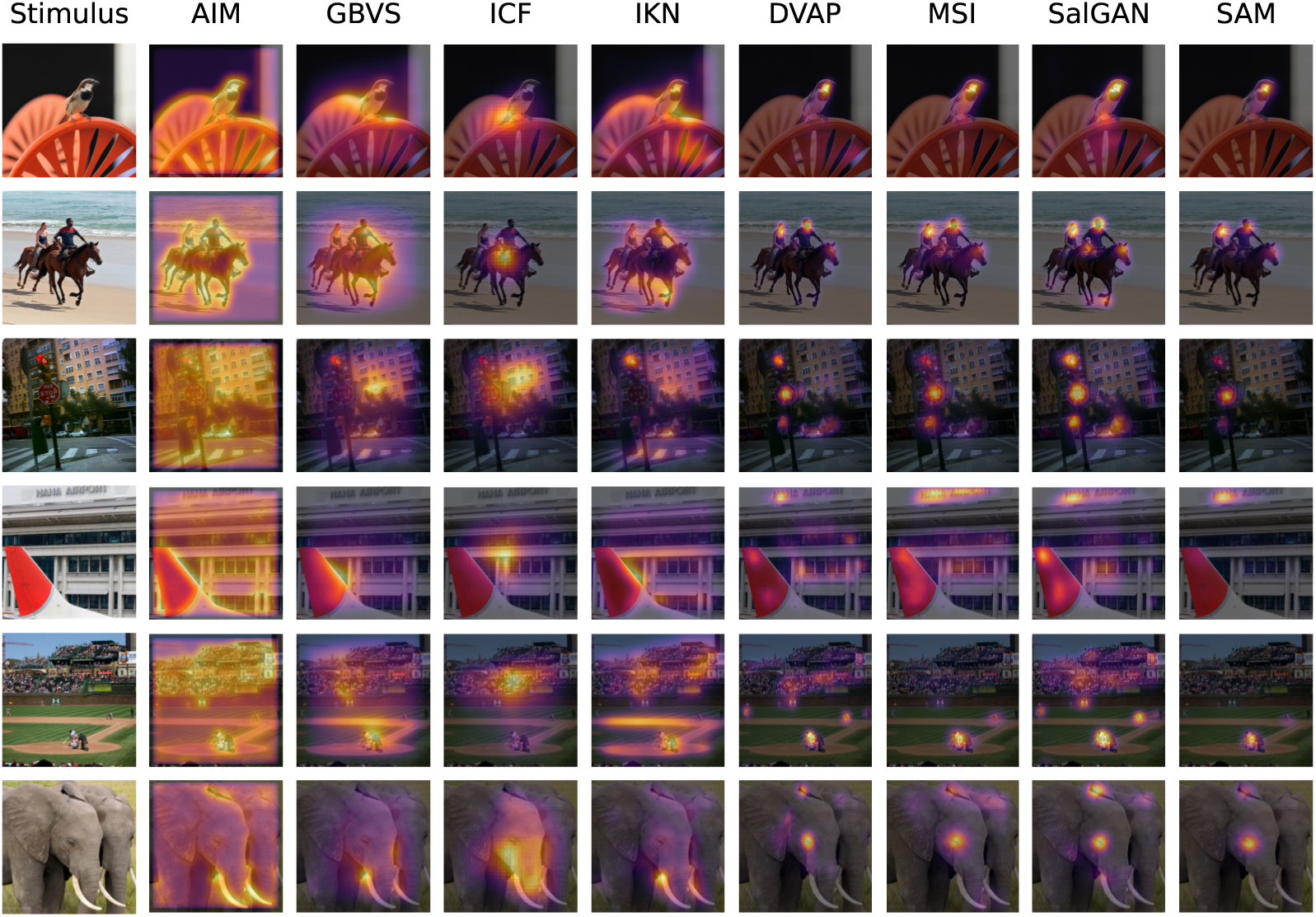
Example stimuli with the predicted saliency maps from low-level (AIM, GBVS, ICF, IKN) and high-level (DVAP, MSI, SalGAN, SAM) models. The latter model category captures semantic information, such as faces in humans and other animals, traffic signs, and text. On the contrary, low-level approaches tend to assign higher values to regions of local contrast and thus predict saliency at object boundaries rather than their center. Overall, the predictions by high-level models appear more consistent than the saliency maps generated by low-level models.

## Appendix D. Center Bias Removal

**Figure D.12:**
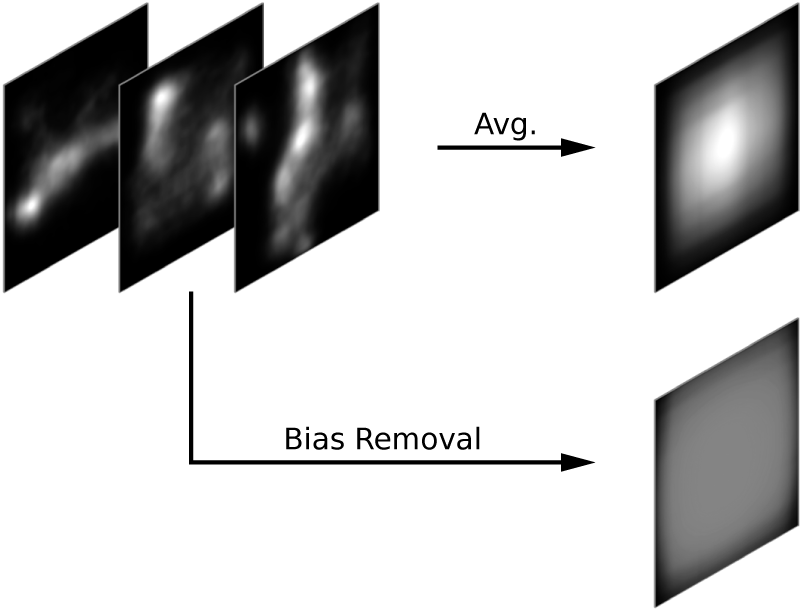
The transformation underlying center bias removal as visualized for saliency maps from the MSI network. A clear center bias emerges after averaging all saliency maps across conditions. When following the two-step procedure described in 2.6, activation patterns become more evenly distributed over the spatial extent of the pre-processed maps. This technique was hence applied to all model and layer outputs.

## Appendix E. Object Segmentation Masks

**Figure E.13:**
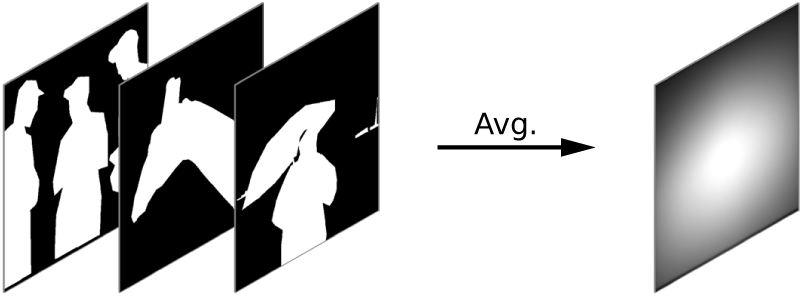
The average segmentation mask across all 73,000 NSD stimuli based on the MSCOCO annotations for a set of 80 object categories. All individual masks were cropped in the same fashion as the corresponding stimuli, pooled per image, and averaged over conditions. The resulting map demonstrates the spatial bias of where objects are presented in the scenes. Evidently, a center preference occurs, which might either be due to the cropping procedure in NSD aimed at minimizing the semantic loss or a general photographer bias [101] in the stimulus set.

## Appendix F. Layer Similarity for Image Classification and Autoencoder Network

**Figure F.14:**
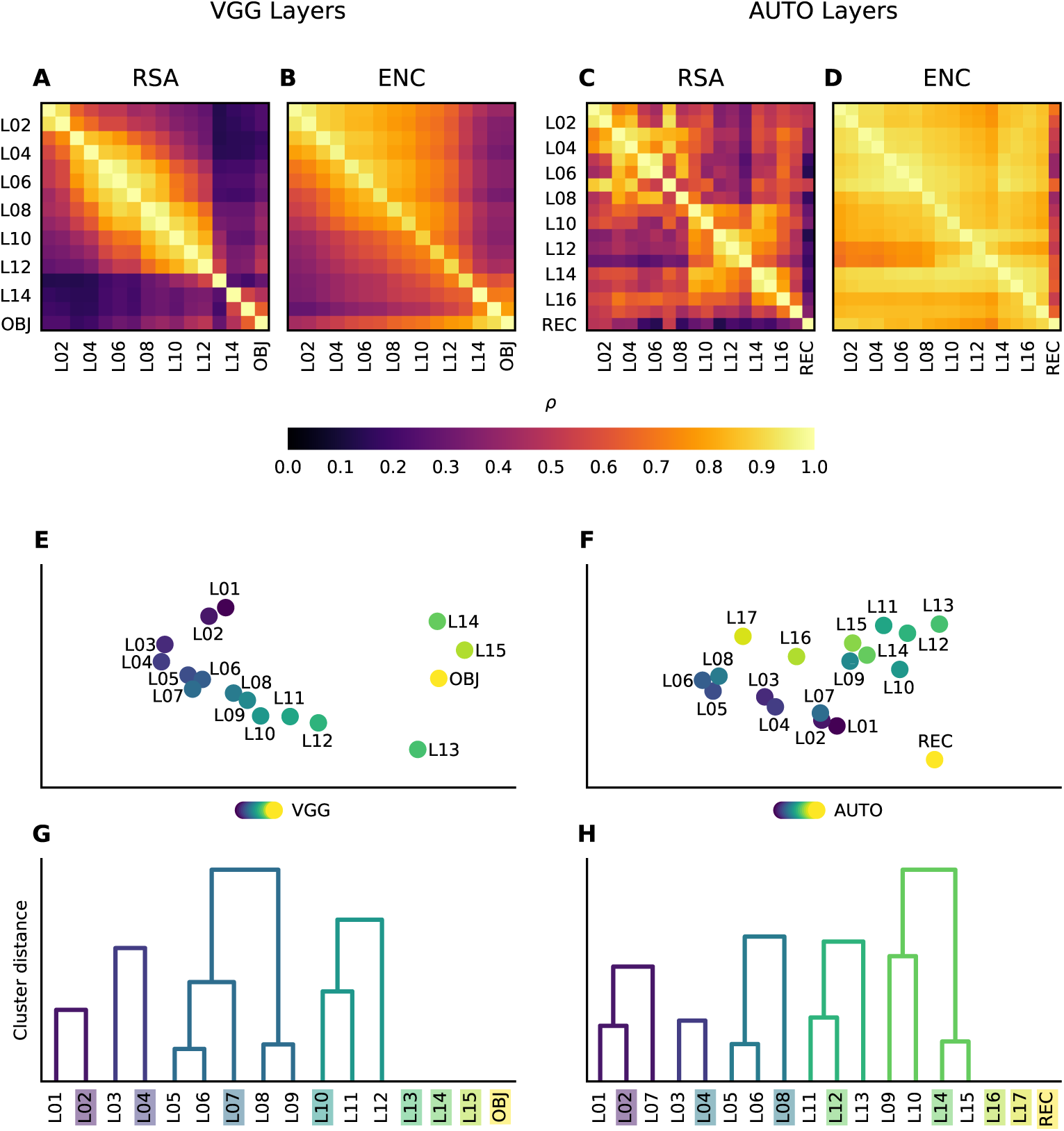
Pairwise comparisons to determine the similarity between layer activations from VGG and an autoencoder network respectively. Both the inputs and desired outputs of the analyses were based on features alone and thus unrelated to neural data. (**A** - **D**) Pearson correlation scores ρ were obtained from two methods: RSA (**A**, **B**) and an encoding model (**C**, **D**). Note that the former yielded symmetrical results along the diagonal whereas the latter introduced a directionality from predictors (x-axis) to targets (y-axis). For ridge regression, elementwise correlation results were averaged across layer activations. (**E** - **H**) An analysis of the previous similarity results obtained from the RSA method. Two techniques were employed to visualize and cluster the pairwise correlation scores: MDS (**E**, **F**) and Agglomerative Clustering (**G**, **H**). The former embeds the similarities between layers in a two-dimensional space whereas the latter forms a dendrogram that represents a hierarchy of clusters. The height at which two clusters merged in the dendrogram indicated the maximum distance between all contained elements. Note that leaf elements without an associated dendrogram formed their own cluster. Colored labels (**G**, **H**) denote which layers were kept for further analyses.

The two methods aimed at evaluating the similarity between model layers revealed that the initial 13 VGG layers strongly resemble the ones from MSI and thus appeared to be mostly unaffected by fine-tuning on the saliency task. The three fully connected layers (L14 to OBJ), and to some degree the last convolutional layer (L13), developed representations distinct from the convolutional stem, since visual features were gradually transformed into class labels. Regarding the autoencoder results, features of the first 13 layers diverged considerably from the VGG weights through fine-tuning on the image reconstruction task. This suggests that the autoencoder benefits less from pre-training compared to the saliency prediction model and formed new representations. Although RSA demonstrated two loose clusters with a boundary halfway through the architecture, correlation results based on the encoding method were consistently high. This could indicate that features were less hierarchically organized, since semantic information is not required for image reconstruction. A notable asymmetry arose for both networks after fitting an elementwise ridge regression. The last layer of each model (i.e. class labels for VGG and reconstructed image for AUTO) was well predicted by the preceding layers, while the reverse did not hold. Similar to MSI, the final representations therefore appeared to be a lossy integration of individual feature channels.

Both clustering techniques confirmed that the three fully connected layers (L14 to OBJ) and the last convolutional layer (L13) of VGG were distinct from the convolutional stem and comparatively unique. The remaining layers neatly clustered with their direct neighbors. By contrast, the autoencoder results demonstrated a looser structure. Clusters also contained non-subsequent layers, which confirms that the representations were less hierarchically organized. Yet the last layer, and to a lesser degree the two previous ones, were again different from the rest. (Figures 11-22)

## Appendix G. Significance of Pairwise Saliency Model Comparisons

**Figure G.15:**
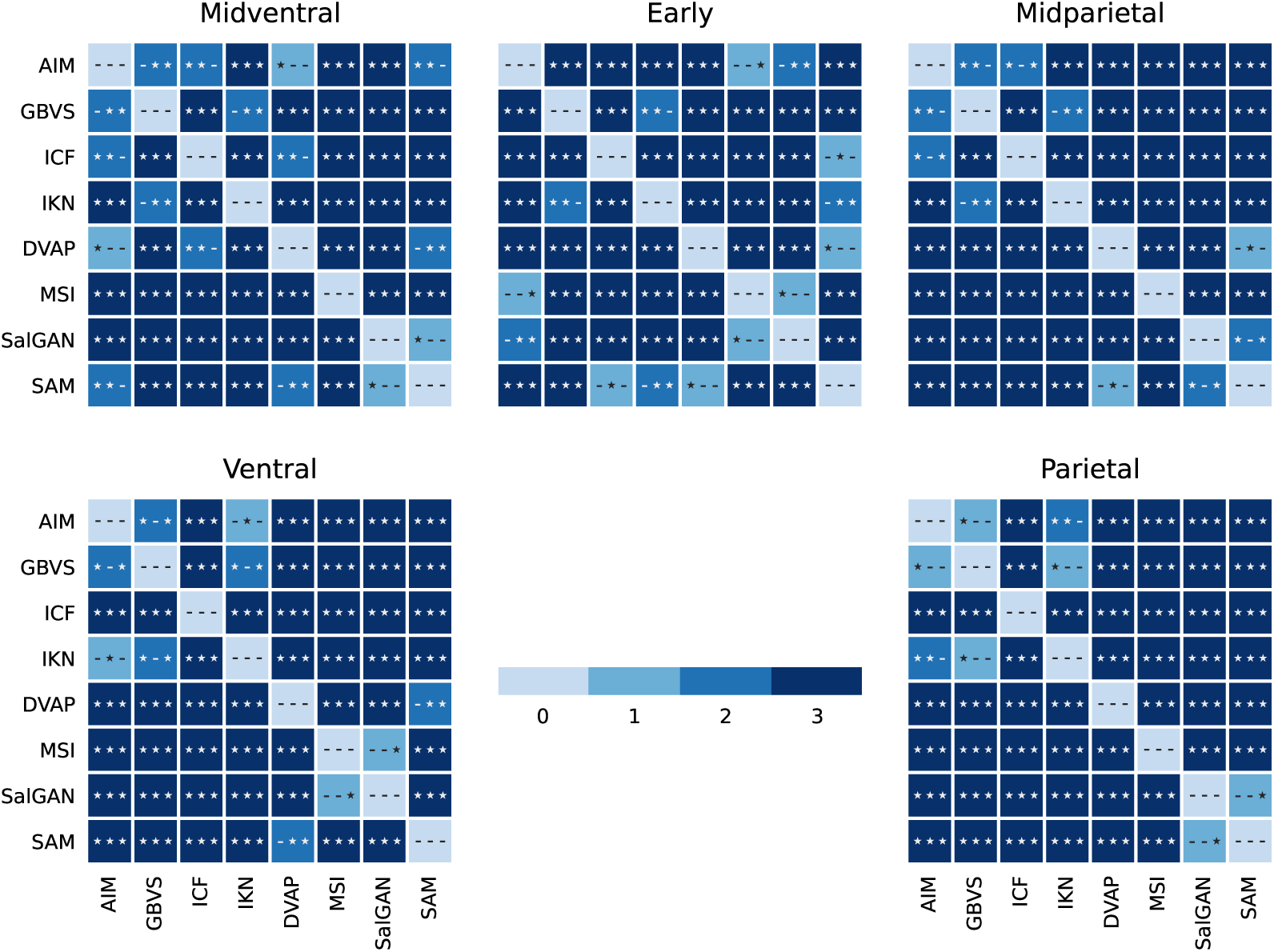
Significance results for the pairwise comparison between saliency models from Figure 4A. All models were individually evaluated and the differences of correlation values were statistically tested (see Section 2.9 for details). The significance was assessed for each method separately in the following order: RSA, ENC, PRF. The symbols (*) and (−) indicate significant and non-significant model differences respectively. Matrices are symmetrical along the diagonal and colors code for the number of significant results across the three methods. For each ROI, the plot shows that most model differences are significant and in agreement for all methods.

## Appendix H. Model Comparison between Saliency and Autoencoder Network

**Figure H.16:**
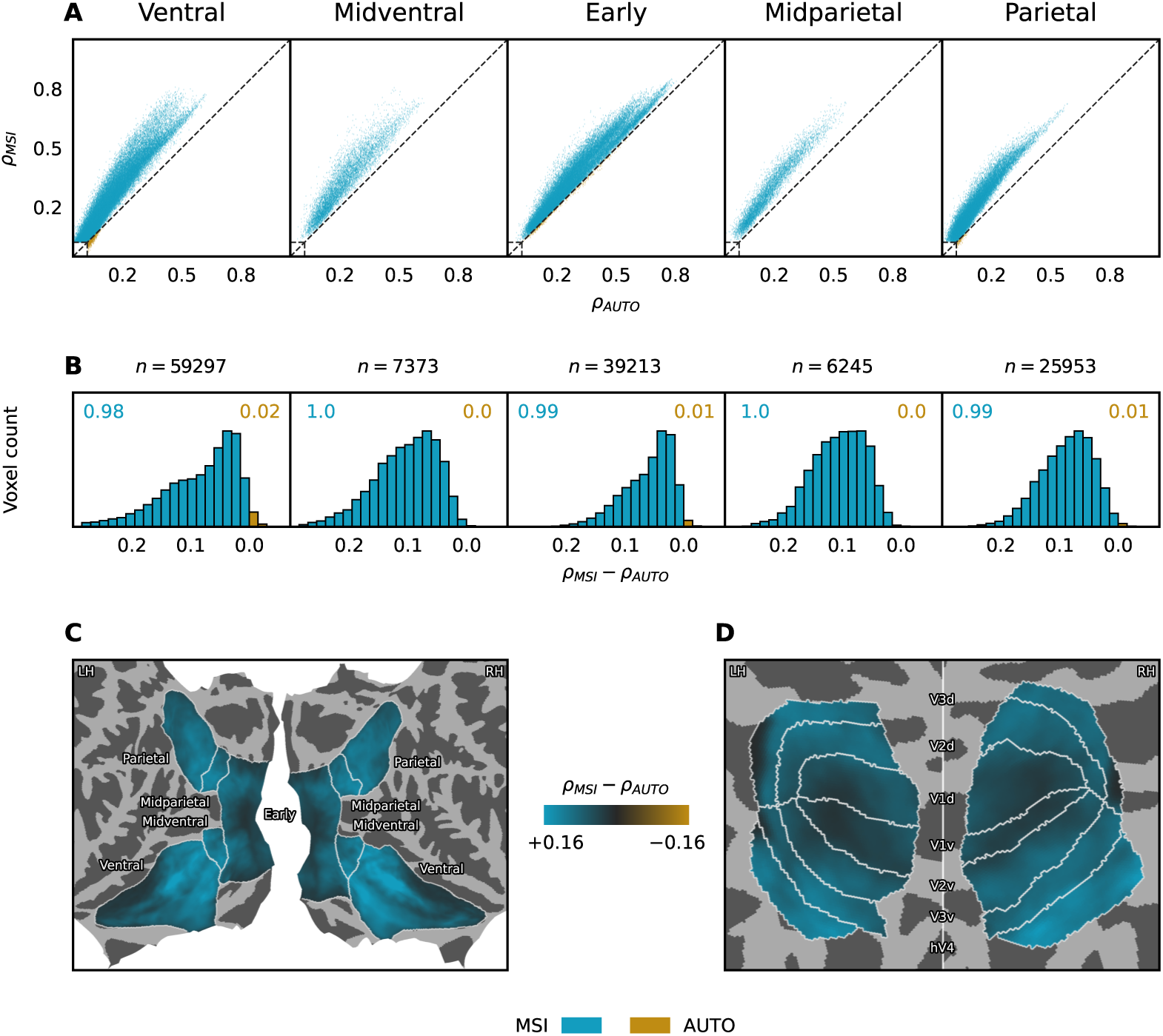
Model comparison between MSI and AUTO with respect to the difference in correlation scores after fitting the two networks independently via ridge regression. (**A**) Voxelwise correlation results for the two models across subjects and per ROI, where points above or below the diagonal indicate an advantage for MSI or AUTO respectively. The rectangles in the bottom left corner of the plots depict the minimum correlation score required for significant results. (**B**) A histogram that aggregates the correlation score differences between the models across all subjects and voxels, with *n* denoting the total number of significant voxels per ROI. The colored numbers at the top corners of the plots summarize the ratio of voxels that were best predicted by each model. All model differences are significant. (**C**, **D**) Group-average correlation differences between MSI and AUTO per vertex on fsaverage. The colormap depicts the magnitude of a model advantage. Voxelwise correlation scores ρ were first obtained per model and subject, projected to a common brain space, and then averaged. Afterwards, the two visual streams (**C**) and retinotopically defined areas (**D**) were visualized on a flatmap and sphere respectively. Note that scores were not normalized by the noise ceiling here.

## Appendix I. Empirical Noise Ceiling Estimates

**Figure I.17:**
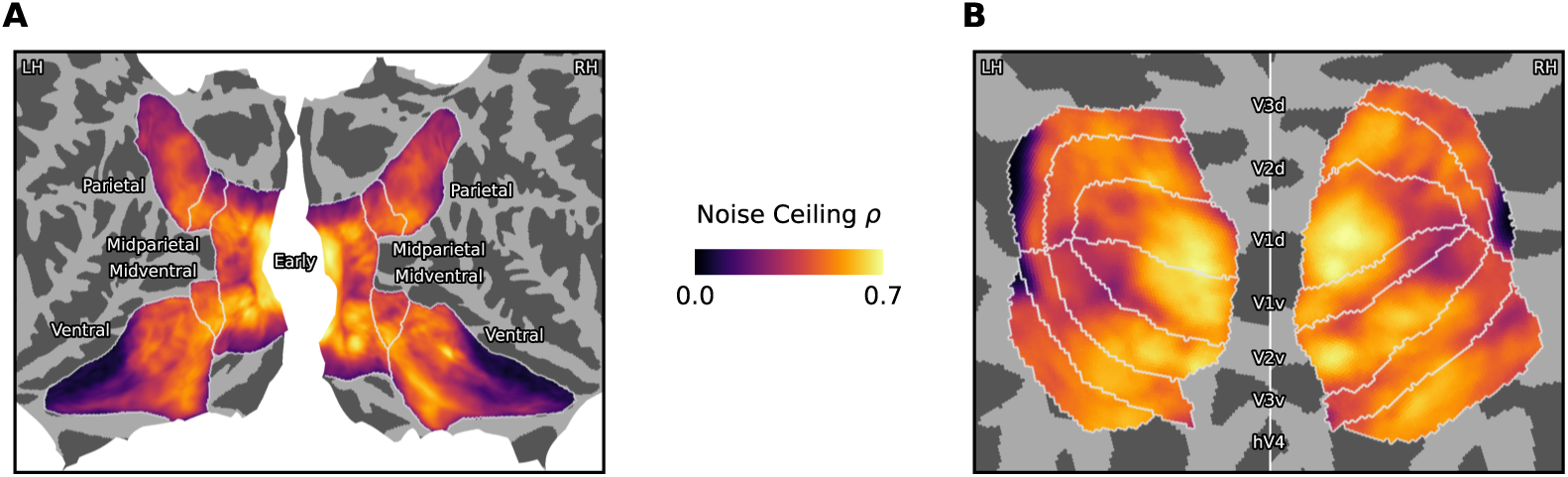
Group-average noise ceiling estimates expressed as the maximum Pearson correlation value ρ per vertex on fsaverage. Voxelwise noise ceiling estimates were first computed per subject (see Section 2.8 for details), projected to a common brain space, and then averaged. Afterwards, the two visual streams (**A**) and retinotopically defined areas (**B**) were visualized on a flatmap and sphere respectively.

## Appendix J. Model Preference and Second-Best Layer Selection

In most cases, the second-best layer of a network occurs immediately before or after the one with the highest performance. One notable exception is V2/V3, where scores of layer L14 were followed by L12 of MSI. The earliest layer under consideration from VGG (L07) achieved the second-best results in vertices along the lingual gyrus that were previously assigned to object representations of OBJ. Finally, neural activations along the fusiform gyrus were well predicted by the most task-specific layer of MSI (L14), as was a small number of vertices located at late processing stages of the parietal ROI. Together with the analysis results from the WTA procedure (see Figures 6C and 6D), it appeared that late layers of MSI were most consistently assigned in midventral areas and parts within ventral regions.

**Figure J.18:**
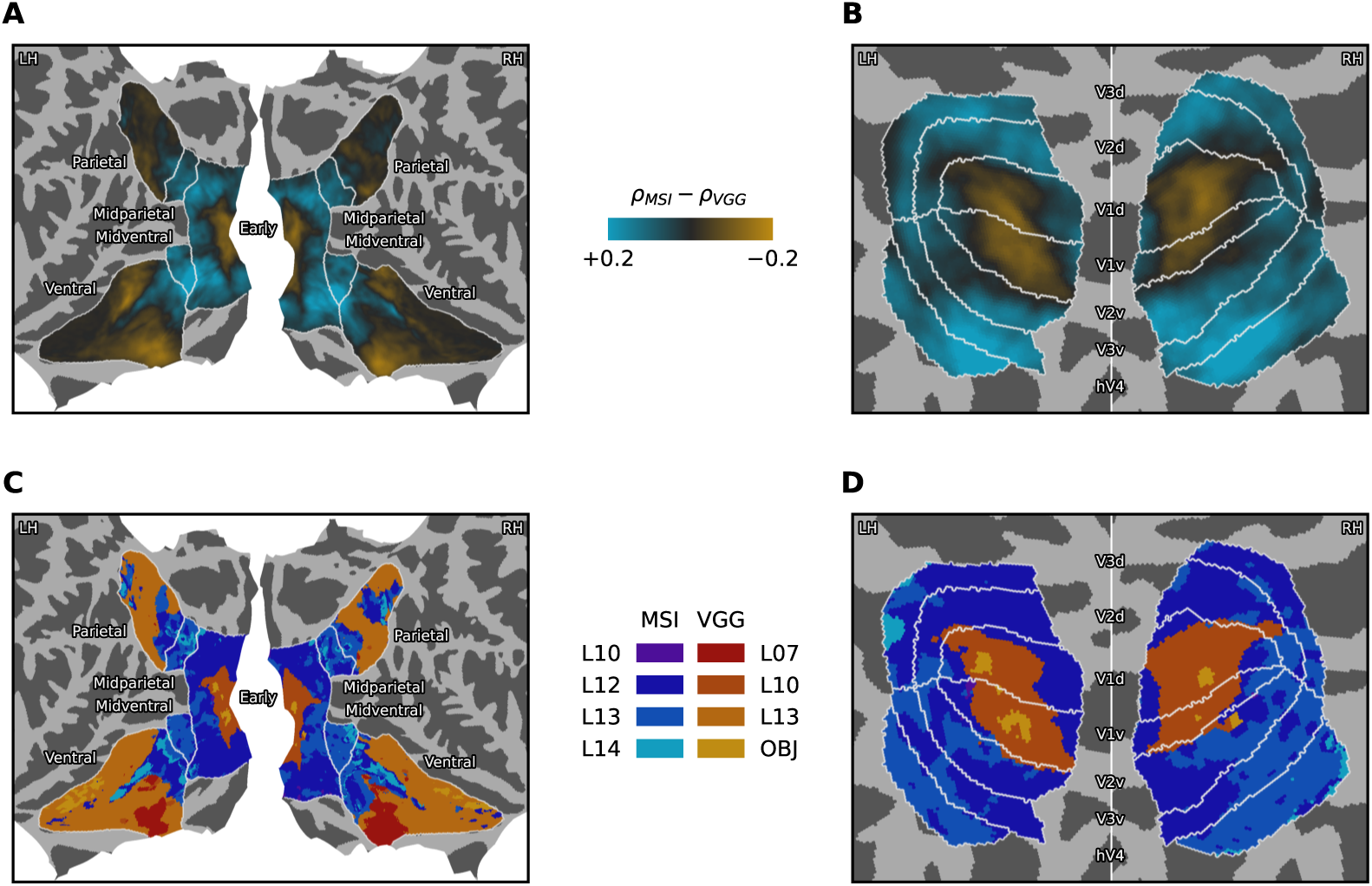
The model and layer preferences after fitting a joint encoding model with eight feature spaces across MSI and VGG. To assign layers to vertices, we first summed the decomposed correlation scores across the four layers from each network, determined the winner, and then chose a layer from the best model with the second-highest individual performance. (**A**, **B**) Group-average correlation differences between MSI and VGG per vertex on fsaverage. The colormap depicts the magnitude of a model advantage after aggregating the contribution of the four layers per network. Voxelwise correlation scores ρ were first obtained per model and subject, projected to a common brain space, and then averaged. Afterwards, the two visual streams (**A**) and retinotopically defined areas (**B**) were visualized on a flatmap and sphere respectively. Note that scores were not normalized by the noise ceiling here. (**C**, **D**) The second-best layer from the selected model was then assigned to each vertex.

## Appendix K. Polar Angle and Eccentricity Estimates from NSD

**Figure K.19:**
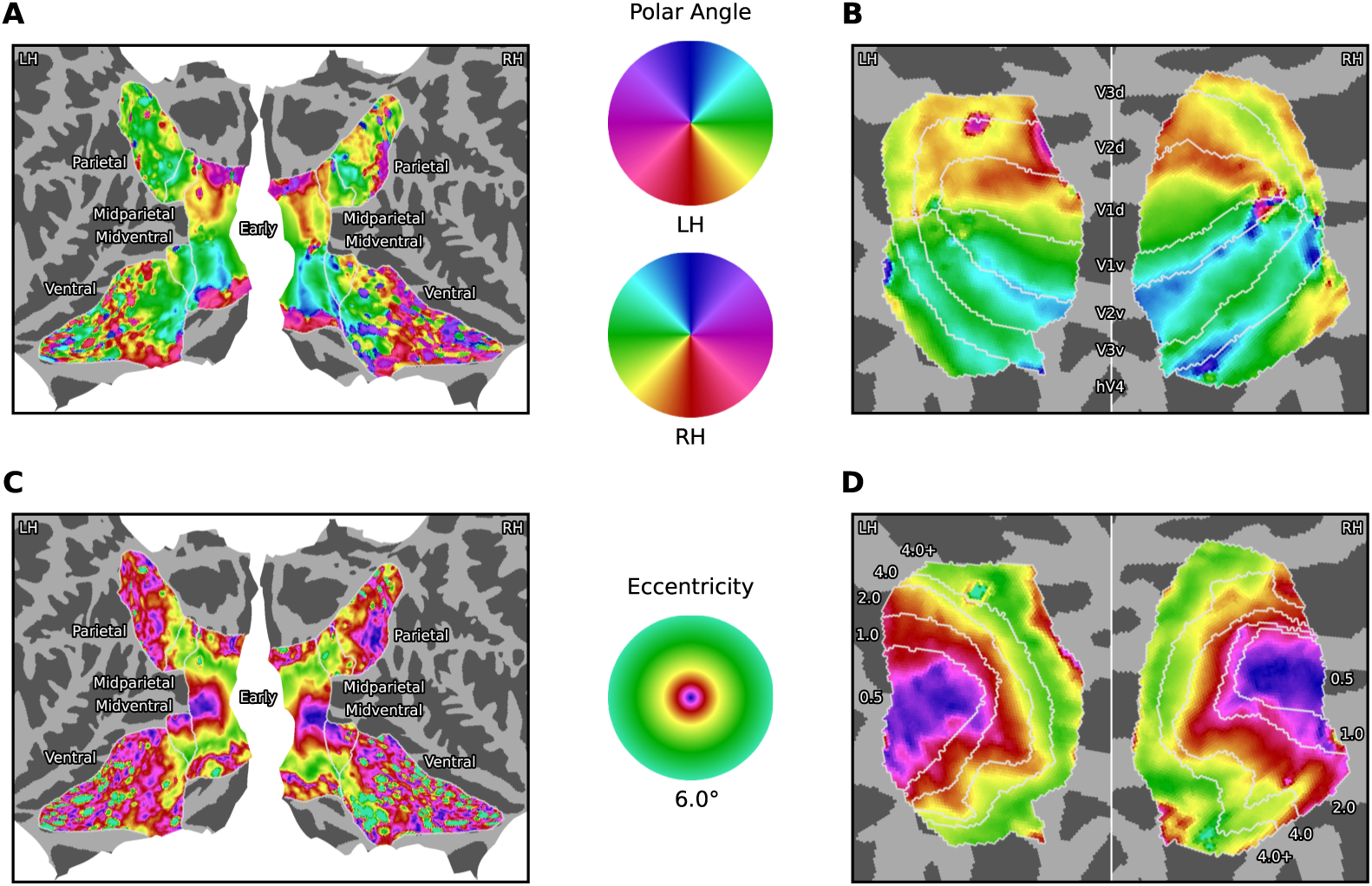
Group-average polar angle and eccentricity estimates from NSD per vertex on fsaverage. Voxelwise receptive field locations were first obtained per subject, projected to a common brain space, averaged, and then converted to polar angle (**A**, **B**) and eccentricity (**C**, **D**) values. Afterwards, the two visual streams (**A**, **C**) and retinotopically defined areas (**B**, **D**) were visualized on a flatmap and sphere respectively. Borders on the sphere visualization for eccentricity estimates (**D**) are based on the eccentricity delineation provided by NSD. The pRF experiment consisted of an ordered presentation of apertures filled with dynamic colorful textures [24].

## Appendix L. Receptive Field Sizes and Fits from MSI Saliency Maps

**Figure L.20:**
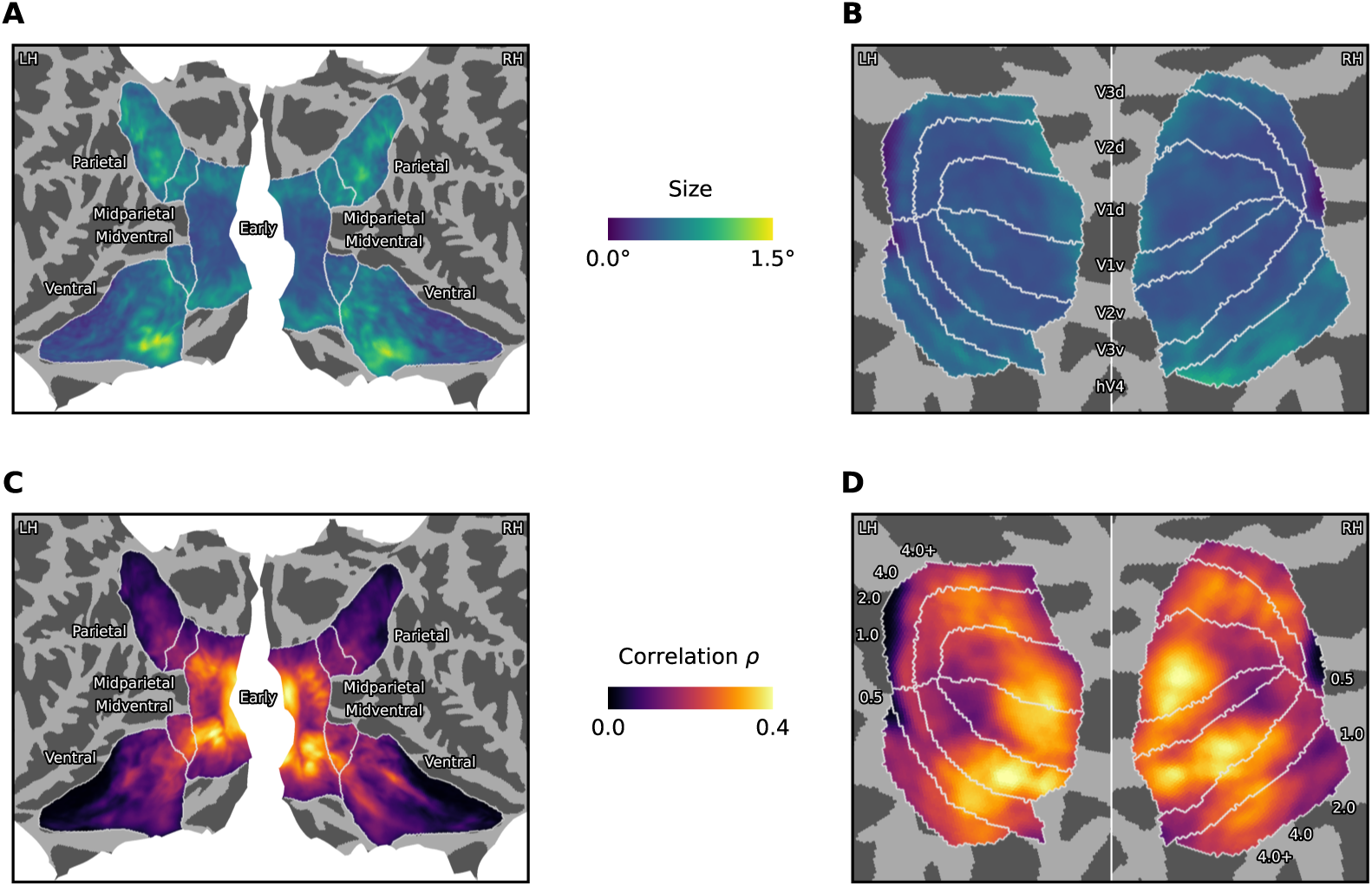
Group-average receptive field size estimates and fits from pRF mapping based on saliency maps of MSI per vertex on fsaverage. Voxelwise receptive field sizes (**A**, **B**) and correlation scores ρ (**C**, **D**) were first obtained per subject, projected to a common brain space, and then averaged. Afterwards, the two visual streams (**A**, **C**) and retinotopically defined areas (**B**, **D**) were visualized on a flatmap and sphere respectively.

## Appendix M. Receptive Field Sizes from MSI Layers

**Figure M.21:**
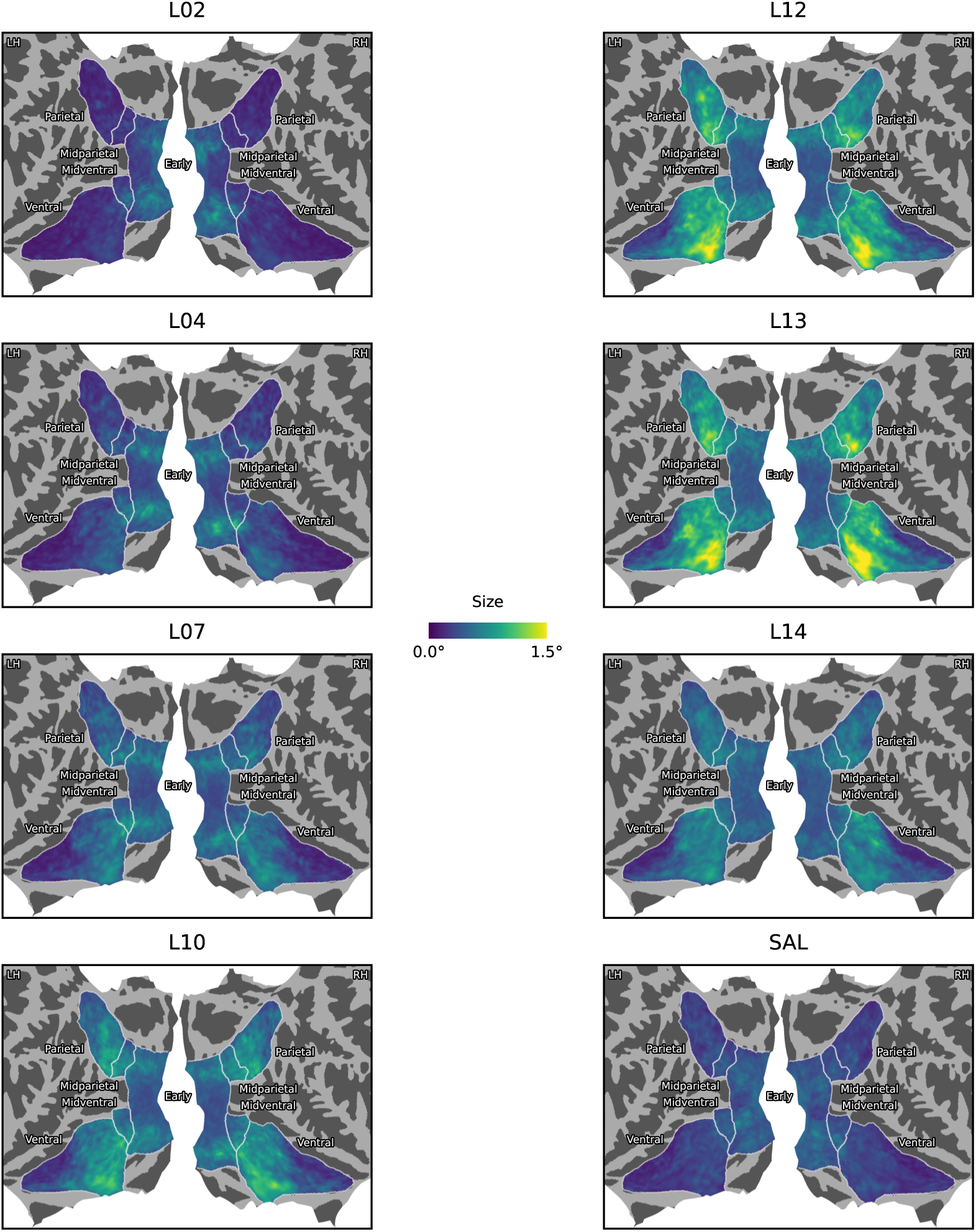
Group-average receptive field size estimates from pRF mapping based on a disentangled subset of eight MSI layers per vertex on fsaverage. Voxelwise receptive field sizes were first obtained per subject, projected to a common brain space, and then averaged. Afterwards, the two visual streams were visualized on a flatmap for each layer separately.

## Appendix N. Receptive Field Fits from MSI Layers

**Figure N.22:**
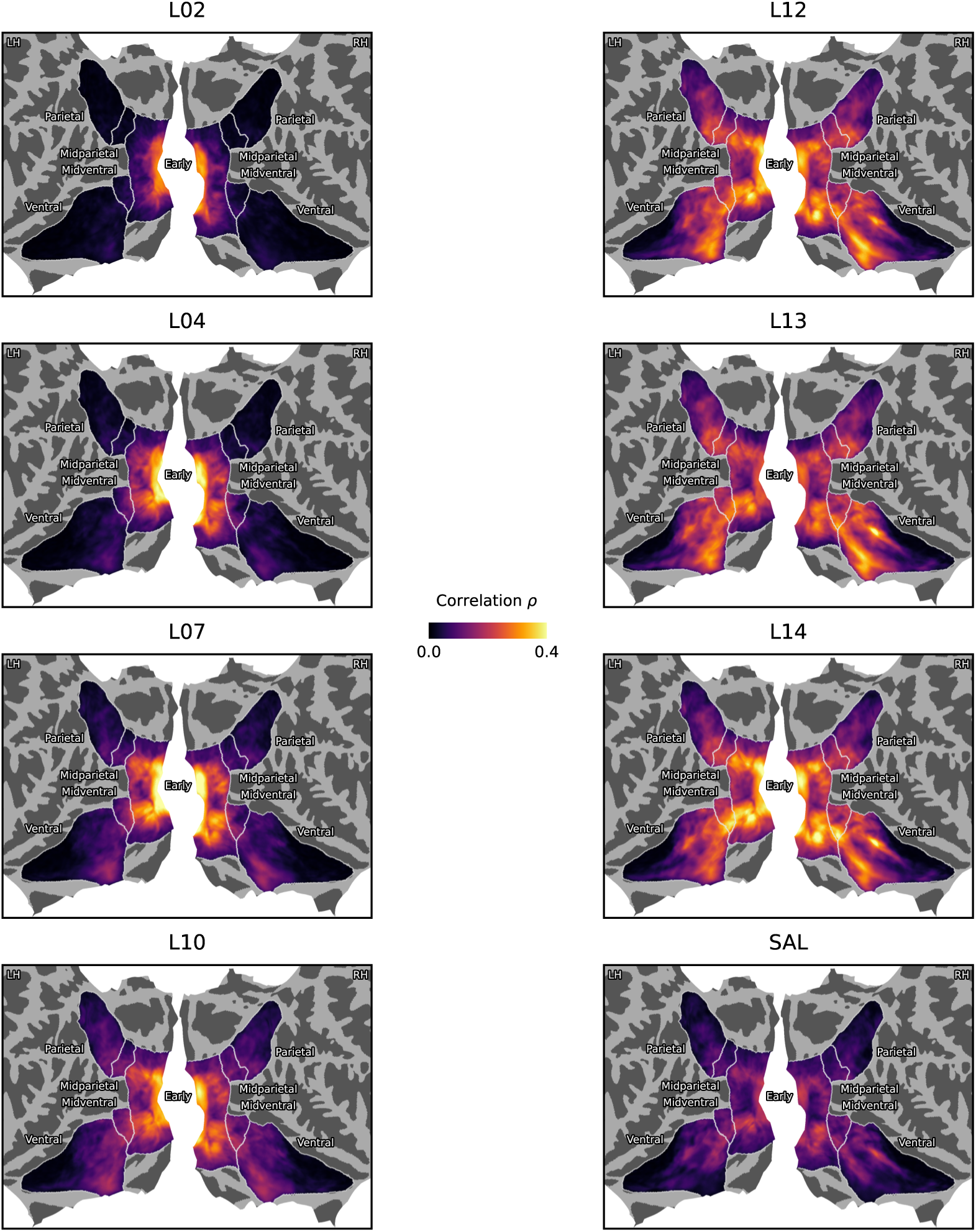
Group-average fits from pRF mapping based on a disentangled subset of eight MSI layers per vertex on fsaverage. Voxelwise correlation scores ρ were first obtained per subject, projected to a common brain space, and then averaged. Afterwards, the two visual streams were visualized on a flatmap for each layer separately.

